# Spontaneous preputial gland infection in *Staphylococcus aureus*-colonized male C57BL/6 mice triggers a Th17-driven immune response

**DOI:** 10.1101/2025.09.05.674392

**Authors:** Liliane Maria Fernandes Hartzig, Shruthi Peringathara, Murty Narayana Darisipudi, Sean Lando Levin Seegert, Elisa Bludau, Stefan Weiss, Antje Vogelgesang, Janosch Schoon, Stefan Gross, Björn Corleis, Barbara M. Bröker, Silva Holtfreter

**Affiliations:** Institute of Immunology, University Medicine Greifswald, 17475 Greifswald, Germany; Department of Functional Genomics, University Medicine Greifswald, 17475 Greifswald, Germany; Clinic and Polyclinic for Neurology, University Medicine Greifswald, 17475 Greifswald, Germany; Center for Orthopaedics, Trauma Surgery and Rehabilitation Medicine, University Medicine Greifswald, 17475 Greifswald, Germany; Department of Internal Medicine B, University Medicine Greifswald, 17475 Greifswald, Germany; Institute of Immunology, Friedrich-Loeffler-Institute, 17493 Greifswald - Isle of Riems, Germany

**Keywords:** *Staphylococcus aureus*, colonization, endogenous, infection, Th, antibodies, preputial gland abscess, abscess, IL-17

## Abstract

Colonization with the pathobiont *Staphylococcus aureus* increases the risk of endogenous *S. aureus* infections if the balance between host and microbe is disturbed. We developed a model of persistent *S. aureus* colonization using the mouse-adapted strain JSNZ. The bacteria transfer from parents to offspring, producing lifelong, usually asymptomatic colonization. Here we report that *S. aureus*-colonized adult male mice frequently develop spontaneous preputial gland infections (preputial gland adenitis, PGA), which are characterized by pronounced pus production and gland enlargement. This study aimed to characterize PGA in terms of phenotype, causative agents, and the pathogen specific antibody and T cell responses. We compared three groups: naïve mice, colonized PGA-negative mice, and colonized PGA-positive mice. PGA occurred in 8/12 (67%) of male breeding animals and in 17/25 (68%) of adult male offspring. The infection did not self-resolve and persisted for several months. Genotyping identified the colonizing strain JSNZ as the causative agent. The infection caused purulent inflammation, with massive bacterial aggregates and neutrophil infiltrates filling the gland lumen. This inflammation completely disrupted the glandular architecture. PGA induced a strong but localized release of IL-1α, IL-1β, IL-17, MIP-1α, and KC in the infected gland. T cells from PGA-draining lymph nodes, as well as splenocytes, reacted to *in vitro* re-stimulation with a *S. aureus* antigen cocktail with the proliferation of Th17 cells, and the release of IL-17 and IFN-γ, corresponding to a type 3/1-immune response. Colonized PGA-positive mice also mounted a robust *S. aureus*–specific serum antibody response. In conclusion, the pathology of this spontaneous, chronic *S. aureus* infection is driven by a strong, type 3–biased immune response that, however, fails to clear the bacteria. This endogenous PGA model provides a valuable tool for studying host– pathogen interactions in natural *S. aureus* infection.

**Author Summary:** Around one in five people naturally carry *Staphylococcus aureus* bacteria in their nose, usually without any problems. But if something upsets the balance, such as a cut in the skin or a weakened immune system, these bacteria can cause infections ranging from boils and abscesses to wound infections and sepsis. To understand how this shift happens, we developed a mouse model in which mice carry *S. aureus* in their nose, much as people do. Over half of the male mice went on to develop abscesses caused by their own colonizing strain. The mice mounted a strong immune response, with local inflammation, a Th17-driven T cell response, and antibodies, yet they did not clear the bacteria. This mirrors the situation in people, in whom *S. aureus* infections are often persistent. The new model lets us study what tips the balance and turns a harmless resident into a cause of disease. By identifying these triggers in future studies, we hope to find better ways to prevent and treat *S. aureus* infections.

## Introduction

*Staphylococcus aureus* (*S. aureus*) remains one of the most lethal pathogens worldwide, causing over a million deaths each year [1]. As a highly adaptable opportunistic bacterium, *S. aureus* colonizes diverse human niches, including the nasal vestibulum, throat, skin, and gastrointestinal tract [2]. Colonization is usually asymptomatic. However, the presence of *S. aureus*-specific antibodies and T cells in human adults suggests that even asymptomatic colonization may involve minor, subclinical breaches of epithelial barriers that trigger systemic adaptive immune responses [3,4]. Moreover, most people also experience self-limiting superficial skin infections with *S. aureus*, such as folliculitis. Furunculosis, cutaneous abscesses, and impetigo are more severe manifestations of *S. aureus* skin and soft tissue infections (SSTIs) [5,6]. SSTIs are among the most common infections encountered in both ambulatory and hospital settings [7]. Their incidence in the United States has increased markedly over the last two decades, driven by the spread of community-acquired methicillinresistant *S. aureus* (CA-MRSA)[7]. SSTIs and their complications are a significant clinical burden, often leading to hospitalization, surgery, bacteremia and, in some cases, even death [5,6]. Beyond SSTI, *S. aureus* can also cause severe, life-threatening systemic conditions such as endocarditis and sepsis [6,8].

Persistent colonization with *S. aureus*, particularly of the skin and nasal passages, is a significant risk factor for the acquisition and recurrence of SSTI [9,10]. Consequently, decolonization strategies employing intranasal mupirocin and chlorhexidine have demonstrated efficacy against such endogenous infections [11,12]. Rising antimicrobial resistance to both agents, however, has prompted the search for alternatives such as povidone-iodine and octenidine [12–15]. Although staphylococcal SSTIs are well established as endogenous infections, the mechanisms by which asymptomatic colonization progresses to invasive disease remain largely elusive [16,17].

Efforts to understand the mechanisms driving the transition from colonization to infection are hampered by the limitations of current animal models. Most SSTI models rely on artificial inoculation, typically by subcutaneous or intradermal injection, and therefore reflect neither the endogenous nature of SSTIs nor the early host-pathogen interactions [18]. In a first attempt, Ge et al. induced a transition from colonization to invasion by tipping the balance between *S. aureus* and the antibacterial immune defense via neutrophil depletion [17]. A disease-specific model of spontaneous SSTI arising from persistent colonization would advance our understanding of disease pathogenesis and facilitate the development of targeted interventions.

CD4^+^ T cells, especially T helper 17 (Th17) and Th1 cells, play a key role in orchestrating a protective immune response against *S. aureus* during colonization and SSTI [4,19,20]. Th17 cells recruit phagocytes and promote barrier function, while Th1 cells enhance the antibacterial effector mechanisms of macrophages [4]. Nasal colonization with *S. aureus* induces a Th1/Th17-dominated T-cell response in both humans and mice [21–24]. Enhanced Th17 activity has been linked to lower *S. aureus* colonization rates in female mice than in males [25]. Similarly, *S. aureus* SSTIs are associated with a local Th17/Th1 response in otherwise healthy individuals [26,27]. For example, Hendriks et al. showed that tissue-resident memory CD4^+^ T cells in human skin respond to *S. aureus* by producing IL-17A, IL-22, IFN-γ, and TNF-α, but not type 2 cytokines, indicating a strong type 1/3 bias [27]. In contrast, Th2-skewed responses have been observed in patients with atopic dermatitis (AD), where *S. aureus*-specific memory CD4^+^ T cells express IL-4 and IL-13, whereas Th17 and Th1 subsets are diminished [28,29].

Our group has established a mouse model of persistent *S. aureus* colonization using the mouseadapted strain JSNZ (clonal complex 88) [30–32]. JSNZ persistently colonizes the nose and gastrointestinal tract and is vertically transmitted to offspring in C57BL/6 colonies [30,33]. While colonization is usually asymptomatic in the breeding pairs and their offspring, we observed spontaneous preputial gland adenitis (PGA) in several older males, but not in females [33]. PGA is a localized inflammatory condition of the preputial glands, paired exocrine glands important for reproductive and behavioral functions [34]. Although it has been sporadically reported in laboratory mice [35], data on the prevalence, age at disease onset, and chronicity of PGA are limited. Moreover, the microbial etiology of PGA, the role of endogenous infection, and immune defense mechanisms involved remain poorly understood. As the colonizing strain JSNZ was originally isolated from a PGA outbreak in a C57BL/6 colony in New Zealand [30], we hypothesized that the observed PGA cases were caused by JSNZ.

Here, we report that spontaneous PGA frequently occurs in male mice persistently colonized with *S. aureus* JSNZ. We characterized its clinical presentation, microbial etiology and pathogen-specific immune response. We identified the colonizing strain JSNZ as the causative agent of the glandular swelling, neutrophil infiltration, and purulent inflammation in infected glands. Immune profiling of re-stimulated lymph node cells and splenocytes revealed a Th17/Th1 response profile. The infection also induced a robust *S. aureus*-specific serum antibody response. This novel model of endogenous infection in *S. aureus* carriers will advance our understanding of host-pathogen interactions in natural *S. aureus* infections.

## Results

### Male mice colonized *with S. aureus* JSNZ frequently develop preputial gland adenitis

Persistent nasal colonization with the mouse-adapted *S. aureus* strain JSNZ is usually asymptomatic in female and young male laboratory mice [30]. However, in our *S. aureus*-colonized C57BL/6NRj colony, we frequently observed preputial gland adenitis (PGA) in male breeding animals and male offspring (Fig 1A). The preputial glands are paired, opaque, flattened structures in the caudal subcutaneous tissue just lateral to the penis base (Fig 1B) [34]. Upon skin removal, markedly enlarged preputial glands containing yellowish purulent material were evident in infected mice (Fig 1C). PGA manifested as unilateral (Fig 1D, n = 9) or bilateral (Fig 1E, n = 9) subcutaneous swellings that progressed to palpable, indurated masses or glandular atrophy (Fig 1F). These swellings were often accompanied by localized skin inflammation, and sometimes ruptured through the skin, leaving scars.

**Fig 1.**
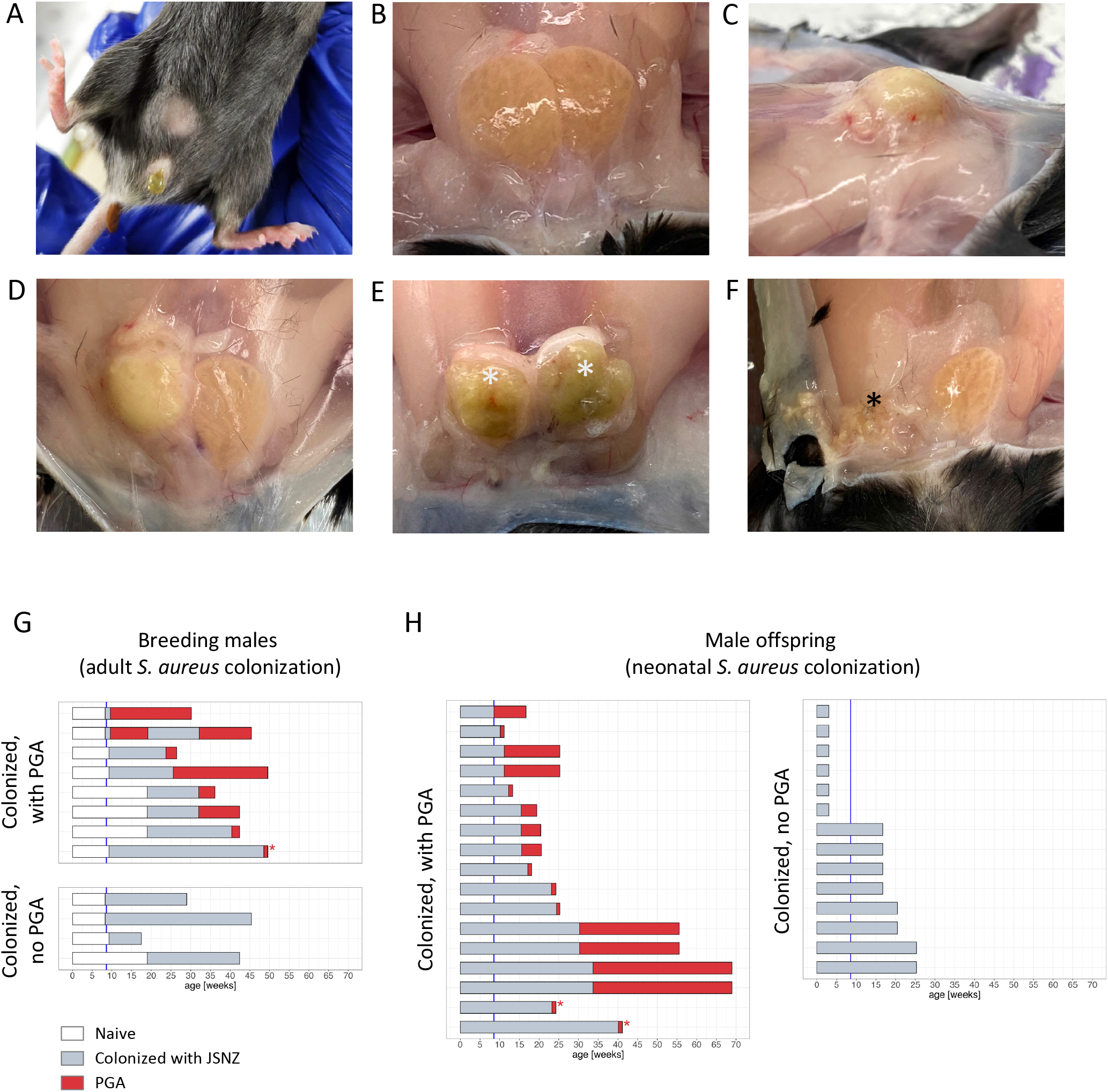
*S. aureus* JSNZ-colonized male mice frequently develop PGA. (A-F) Macroscopic presentation of healthy and infected preputial glands in male C57BL/6NRj mice. (A) PGA-related unilateral subcutaneous swelling in the lower abdomen, (B) paired preputial glands in healthy male mice, (C, D) unilateral PGA with pronounced ectasia, (E) bilateral PGA (white asterisk), and (F) late-stage unilateral PGA with shrunken glands (black asterisk). (G-H) Time course of PGA development in male JSNZ-colonized mice: (G) breeding males, (H) male offspring. In three mice the onset of PGA was not documented (red asterisk). White bars: *S. aureus*-free, grey bars: JSNZ-colonized, no PGA; red bars: JSNZ-colonized, PGA, blue line: onset of fertility (8 weeks).

To assess the prevalence, age at onset, and chronicity, we monitored PGA in our documented cases (Fig 1G, H). PGA occurred in 8/12 (67%) breeding males and 17/25 (68%) adult male offspring. In breeding males, it developed on average 13.0 weeks (median; range: 1.3 – 21.4 weeks) post JSNZ inoculation. Offspring became colonized shortly after birth [33], yet infection appeared no earlier than 8.6 weeks of age (median: 15.6 weeks, range 8.6 – 33.7 weeks), coinciding with fertility onset.

Clinically, PGA presented as localized swelling and inflammation, progressing to abscesses in severe cases. The condition was generally not self-resolving; once established, symptoms typically persisted until animals were euthanized at experimental endpoints or for welfare reasons after gland rupture. One breeding males showed macroscopic remission after few weeks, though we cannot exclude atrophy of the affected gland with subsequent infection of the adjacent so far unaffected gland. Overall, these findings suggest that PGA is a highly prevalent and persistent condition in JSNZ-colonized male C57BL/6 mice, emerging only after sexual maturity.

### The colonizing JSNZ strain is the causative agent of PGA

To elucidate the PGA pathophysiology, we analyzed three groups: *S. aureus* JSNZ-colonized mice with PGA (n = 19) or without glandular infection (No PGA, n = 9), and *S. aureus*-free naïve controls (n = 11). Microbiological analysis of homogenized glands identified *S. aureus* as the sole pathogen in 18 of 19 PGA cases (Fig 2A). The remaining breeding male with bilateral PGA harbored *Klebsiella pneumoniae* in the left and *S. aureus* in the right gland. Moreover, *Mammaliicoccus sciuri*, an opportunistic pathogen akin to *S. aureus*, was isolated from healthy preputial glands of two naïve mice. *S. aureus*-colonized mice with and without PGA carried high bacterial loads in the nose, cecum, and feces, reflecting persistent colonization of the nose and gastrointestinal tract (Fig 2B-D). *S. aureus* was abundant in PGA glands but absent from those of symptom-free colonized and naïve mice (Fig 2E). No systemic dissemination was detected, corroborating PGA as a localized infection (S1 Fig). Genotyping revealed that the invasive isolates shared the *spa* type (t729) of the colonizing JSNZ strain, identifying PGA as an endogenous infection. Together, the microbiological data show that the colonizing *S. aureus* strain JSNZ is the etiological agent of PGA.

**Fig 2.**
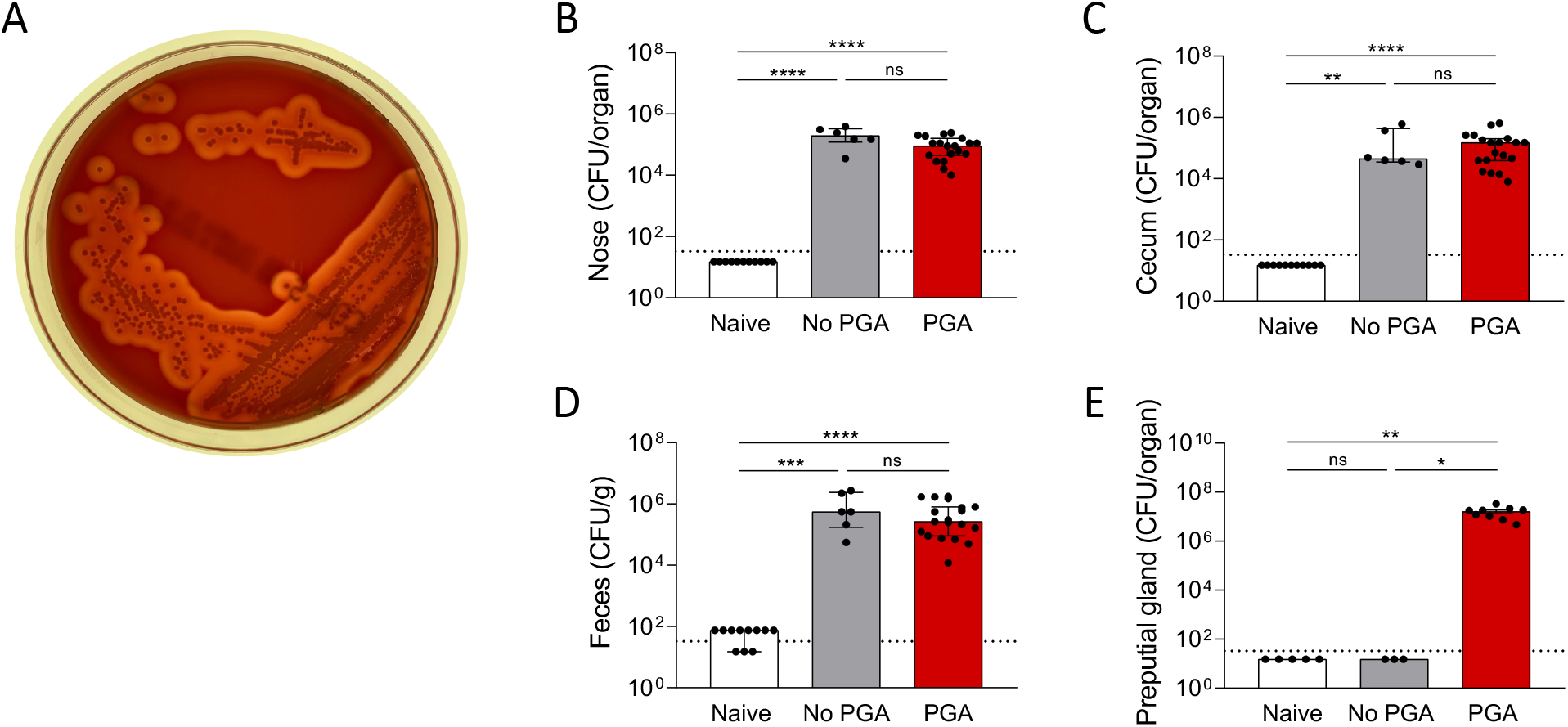
*S. aureus* JSNZ is the causative agent of PGA. (A) Blood agar smear of homogenized PGA tissue showing *S. aureus* in pure culture. (B-E) *S. aureus* CFUs in the nose (B), cecum (C), feces (D), and preputial glands (E). Groups: naïve (*S. aureus*-free); no PGA (JSNZ-colonized, no PGA); PGA (*S. aureus* JSNZ-colonized, with PGA). Each data point represents one animal (median of three technical replicates). Median with interquartile range. Statistics: Kruskal–Wallis test with Dunn’s multiple comparisons. p ≤ 0.05 (*), p ≤ 0.01 (**), p ≤ 0.001 (***), p ≤ 0.0001 (****); ns, not significant.

### PGA is characterized by enlarged glands and massive infiltration of neutrophils

To characterize the pathological changes in PGA, we performed histopathology (H&E) and immunohistochemistry (IHC) on paraffin-embedded gland sections from healthy mice (naïve and colonized without PGA) and infected mice (colonized with PGA). By H&E, healthy glands appeared as well-organized, lobulated structures with prominent excretory ducts (Fig 3, upper left and middle, 4X), containing densely packed sebaceous secretory cells in acini and clear lumina free of infiltrates (Fig 3, lower left and middle, 20X). In contrast, glands from PGA mice showed marked ductal ectasia with multiple dilated ducts and loss of lobular architecture (Fig 3, upper right). At 20X, sebaceous cells were absent and large cellular infiltrates were evident.

**Fig 3.**
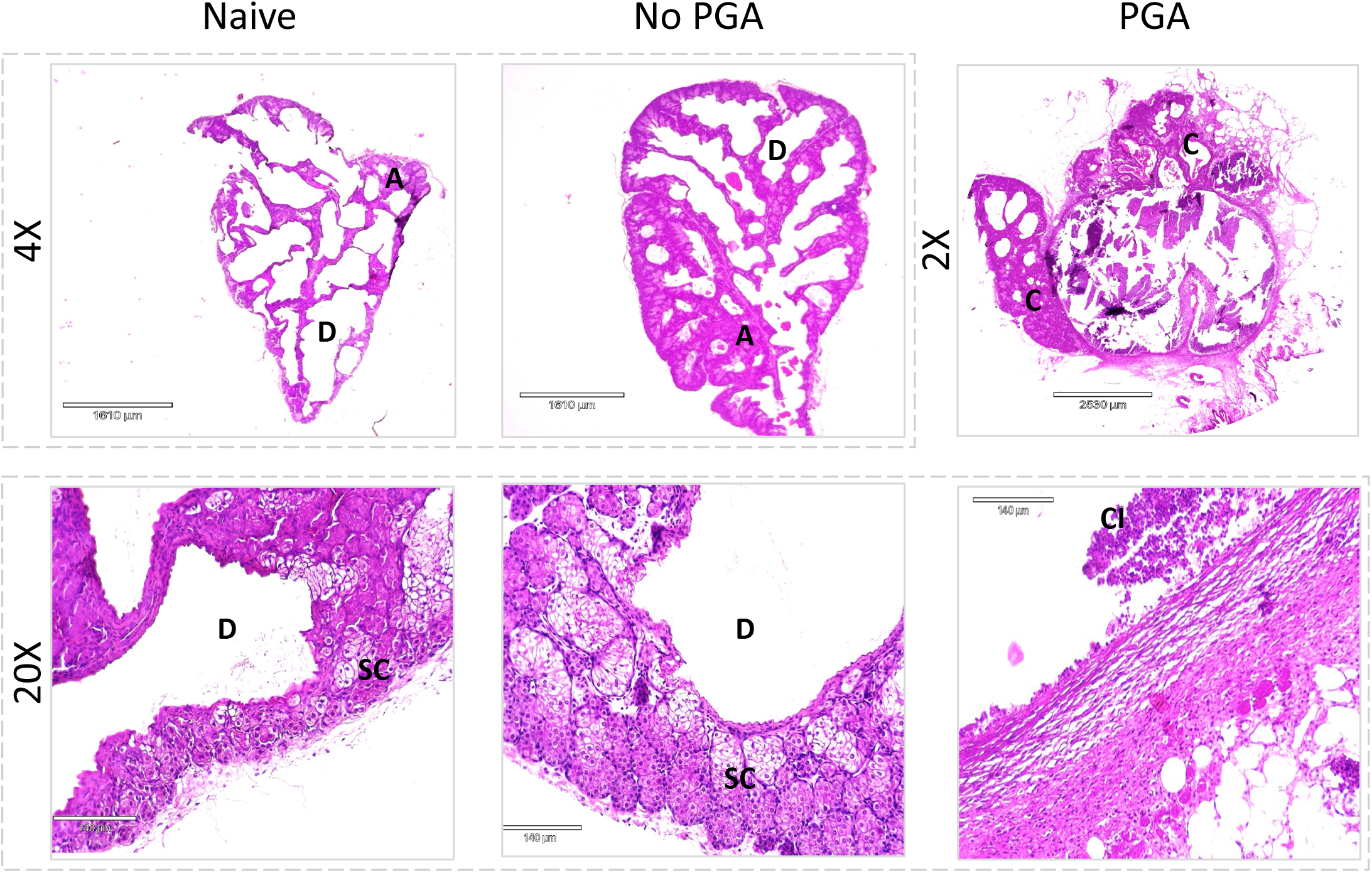
*S. aureus*-induced PGA disrupts glandular architecture. H&E-stained sections of preputial glands from *S. aureus*-free mice (Naïve), JSNZ-colonized mice without PGA (No PGA), and JSNZ-colonized mice with PGA (PGA). The lower magnification provides an overview of the structure of the preputial glands (upper panel). At low magnification (4X, 2X), healthy glands exhibited lobulated morphology comprising ducts (D) and acini (A). In PGA, the architecture was severely distorted, and the lumen was filled with a loosely attached cellular mass. The inflamed gland was often adherent to neighboring connective tissue (C). At high magnification (20X), healthy glands revealed cavernous excretory ducts (D) and sebaceous secretory cells (SC) forming acini (A). PGA specimens showed dilated ducts lined with desquamated epithelium, and large cellular infiltrates (CI). These infiltrates sometimes detached in large portions during cryosectioning and staining. Representative images of naïve (n = 6), No PGA (n = 3), and PGA groups (n = 5). Scale bar, 140 µm.

Pus consists mainly of dead neutrophils recruited to the infection site together with a protein-rich exudate [36]. To define the composition of the purulent mass, we performed IHC staining. As expected, control glands (naïve, No PGA) showed no *S. aureus* or neutrophil infiltration, whereas the lumina of infected glands contained large *S. aureus* clusters often co-localized with massive neutrophil infiltrates (Fig 4).

**Fig 4.**
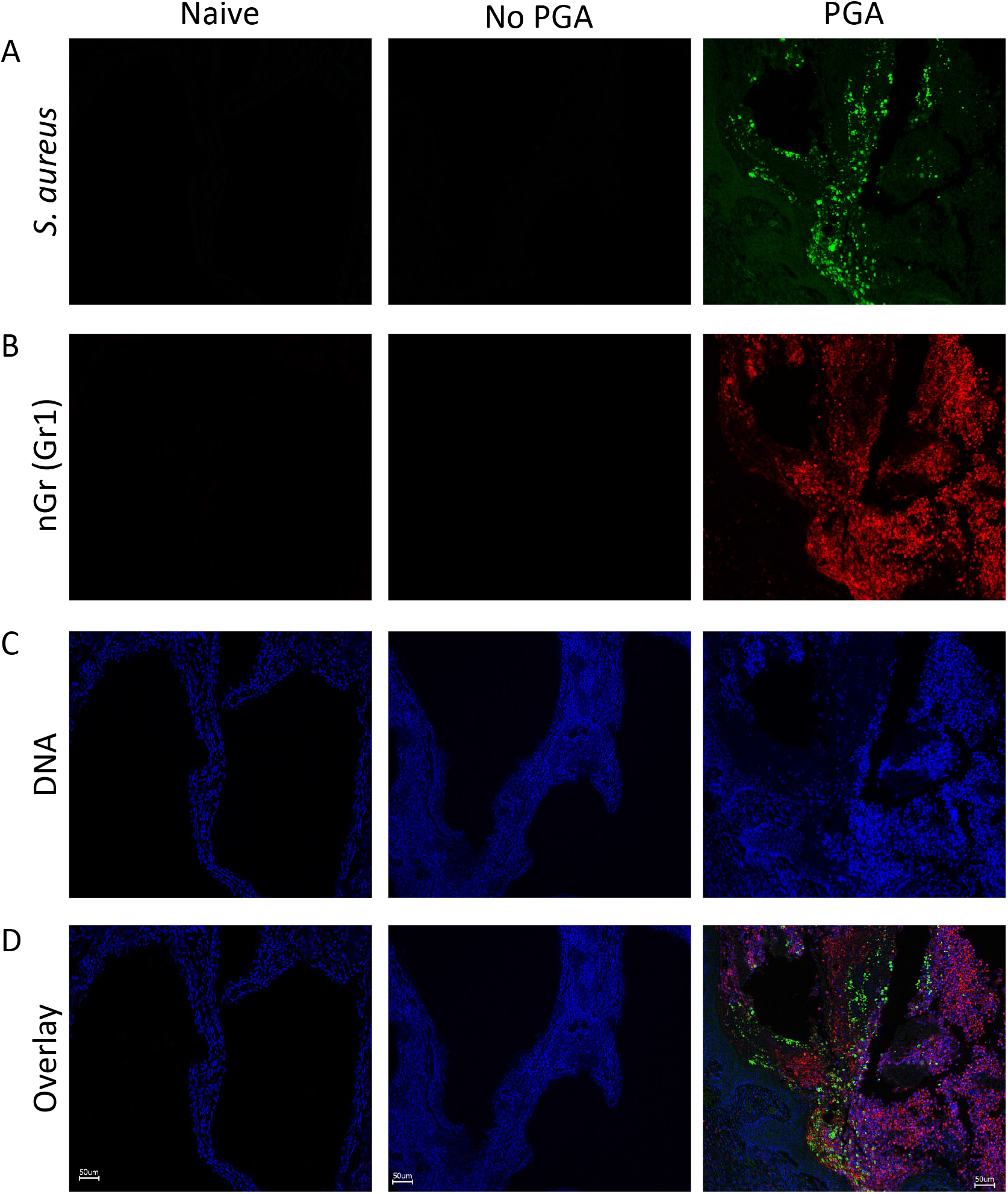
Suppurative inflammation in PGA is characterized by neutrophilic infiltration and *S. aureus* aggregates. Immunohistochemistry of preputial gland sections (4⍰μm), stained for *S. aureus* (anti-*S. a*., green, A), neutrophils (anti-GR1, red, B), and DNA (DAPI, blue, C). Overlay shown in (D). Infected preputial glands exhibited substantial *S. aureus* aggregates with extensive neutrophil infiltration, consistent with purulent inflammation. Healthy glands from naïve and *S. aureus* colonized mice showed no cellular infiltrates. Images are maximum intensity projections of Z-stacks. Representative images of naïve (n = 6), No PGA (n = 3), and PGA groups (n = 5). Scale bar, 50 µm.

In addition, we observed strong MPO activity, which reflects neutrophil activation, in infected glands as compared to healthy glands from naïve or colonized mice (Fig 5A). In cases of unilateral PGA, MPO activity was detected in the inflamed gland, but remained at baseline in the neighboring healthy gland (Fig 5B). Collectively, these findings show that PGA is characterized by loss of glandular integrity with neutrophil infiltration and pus accumulation.

**Fig 5.**
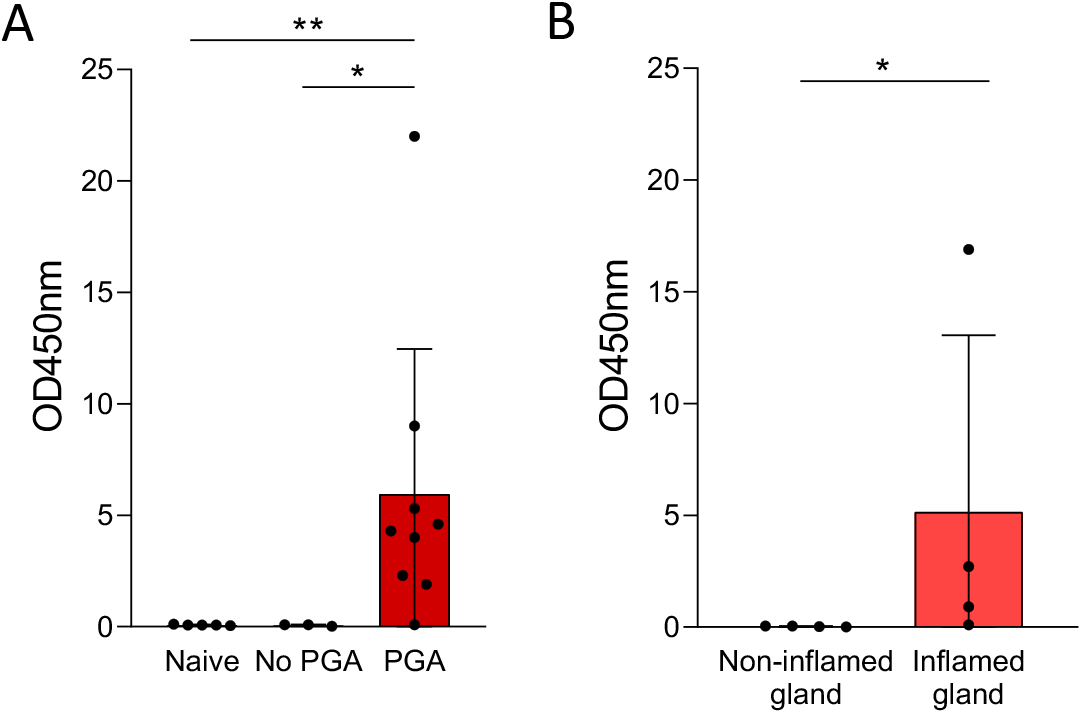
Strong MPO activity in PGA. Peroxidase activity was quantified in preputial gland homogenates using a TMB substrate. (A) Glands from *S. aureus*-free male mice (Naïve, n⍰=⍰5), JSNZ-colonized male mice without PGA (No PGA, n = 3), and JSNZ-colonized mice with PGA (PGA, n⍰=⍰9). (B) Inflamed and non-inflamed glands from 4 mice with unilateral infections. Mean ± SD. Statistics: Kruskal-Wallis test with Dunn’s multiple comparisons (A), Mann-Whitney test (B). p ≤ 0.05 (*), p ≤ 0.01 (**).

### Spontaneous PGA induces a strong local pro-inflammatory chemokine and cytokine response

To investigate the local immune response associated with PGA, we analyzed cytokine and chemokine levels in preputial glands from three groups of male mice: PGA (n = 9), No PGA (n = 3), and naïve controls (n = 5). Paired glands from each mouse were homogenized. In contrast to uninfected glands (naïve and No PGA group), infected glands showed a substantial increase in innate pro-inflammatory cytokines, including IL-6, IL-1α, and IL-1β (Fig 6, left panel). Type 3 cytokines (IL-17A, IL-17F, IL-22) were also strongly elevated, and type 1 cytokines (IFN-γ, TNF-α) increased moderately. In contrast, type 2 cytokines (IL-4, IL-5, IL-13) and the anti-inflammatory cytokine IL-10 were unaltered. Infected glands also contained elevated chemokines, including B lymphocyte chemokine (BLC/CXCL13), macrophage inflammatory protein-1α (MIP-1α/CCL3), and the potent neutrophil chemoattractant KC (keratinocyte-derived chemokine) (CXCL1).

**Fig 6.**
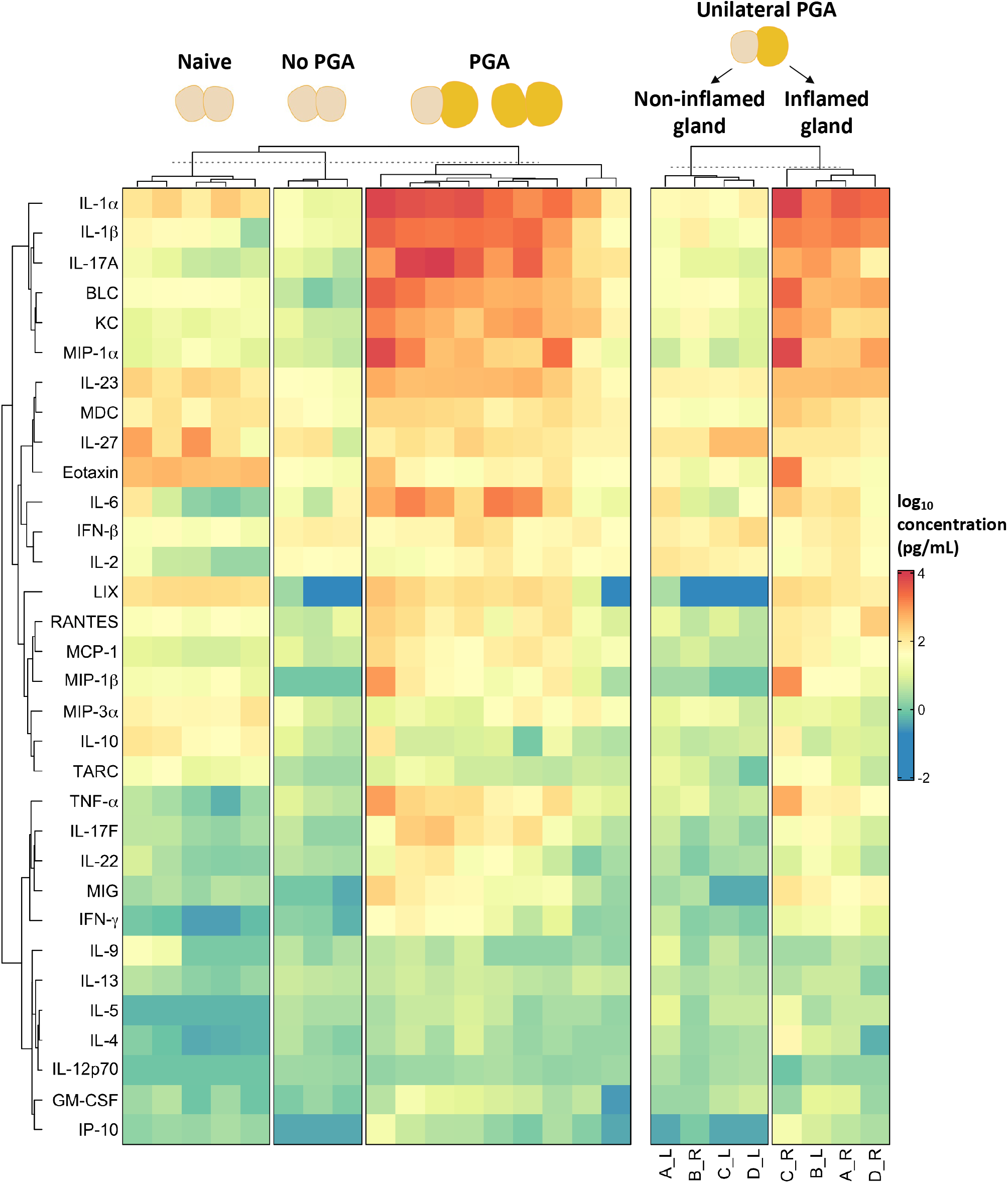
Cytokine and chemokine profiling reveals localized inflammation in PGA. Left panel: Heatmap of cytokine and chemokine concentrations in preputial gland homogenates from *S. aureus*-free male mice (Naïve, n⍰=⍰5), JSNZ-colonized male mice without PGA (No PGA, n = 3), and JSNZ-colonized mice with PGA (PGA, n⍰=⍰9). Paired glands were pooled per mouse. In a subset of mice with unilateral PGA, inflamed and non-inflamed glands from the same animals (n = 4) were analyzed separately (right panel). L, left gland; R, right gland. Concentrations were log_10_-transformed and hierarchically clustered.

As most PGA cases were unilateral, we next assessed whether the inflammatory response was restricted to the infected gland (n = 4). Indeed, the response was again strongly elevated in the diseased gland but remained at baseline in the neighboring gland (Fig 6, right panel). Systemic levels also remained at baseline (S2 Fig). In summary, PGA elicits a massive but locally confined release of pro-inflammatory cytokines and chemokines.

### PGA triggers a *S. aureus*-specific Th17-driven response in the spleen

To characterize the *S. aureus*-specific T cell response in PGA, splenocytes from mice with PGA and from control mice were left untreated or re-stimulated *in vitro* for 4 days with an *S. aureus* antigen cocktail (heat-inactivated extracellular proteins and UV-inactivated cells of JSNZΔ*spa*), which reactivates and expands *S. aureus*-specific T cells. T cell subpopulations were then analyzed by flow cytometry-based phenotyping and by quantifying cytokines in culture supernatants. The control mice, provided by our local animal facility, harbored a CC15 MSSA isolate (spa type t084) in their nose; they were thus not naïve to *S. aureus* but suitable as control group.

Flow cytometry of re-stimulated splenocytes pointed to *S. aureus*-specific Th17 cells in mice with PGA. Re-stimulation raised the percentage of proliferating CD4^+^ T cells (CD4^+^CFSE^low^; not significant), predominantly driven by an increase in Th17 cells (CD4^+^CFSE^low^RORγt^+^) (Fig 7). This expansion was also reflected by a higher overall Th17 proportion among CD4+ T cells. The percentage and absolute number (S3 Fig) of effector/effector-memory (Teff/Tem) Th17 cells (CD4^+^CFSE^low^RORγt^+^CD44^+^CD62L^-^) increased significantly upon re-stimulation of splenocytes from PGA-positive mice. Control mice (CC15-colonized) showed only a moderate, non-significant increase in proliferating CD4+ T cells with the JSNZ antigen cocktail, but no Th17 expansion.

**Fig 7.**
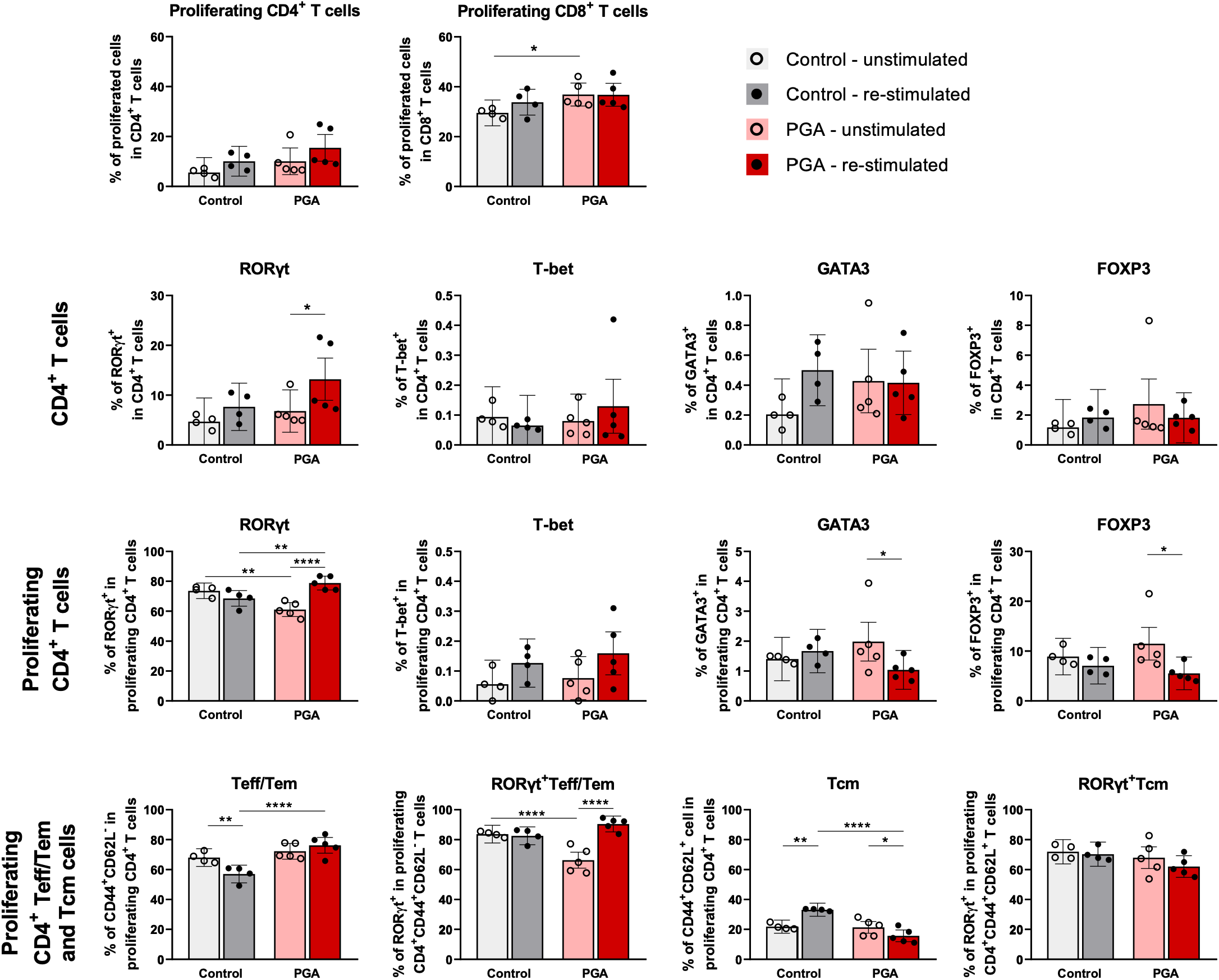
PGA induces a systemic *S. aureus*-specific Th17-driven immune response. Splenocytes from JSNZ-colonized mice with PGA were CFSE-labelled and cultured with or without an *S. aureus* antigen cocktail for 4 days. Mice initially considered *S. aureus*-naïve but retrospectively found to carry a CC15 isolate served as the colonized PGA-negative control. CD4^+^ and CD8^+^ T cell proliferation and the frequency of lineage-specific transcription factors RORγt, T-bet, GATA3 and FOXP3 within Th cell populations were assessed by flow cytometry. Within CD4^+^ T cells, effector/effector memory T cells (Teff/Tem, CD44^+^CD62L^-^) and central memory T cells (Tcm, CD44^+^CD62L^+^) subsets were identified. n = 4-5 mice/group. Mean ± SI. Statistics: linear mixed model (including animal-ID as random factor) in two-way ANOVA design. p ≤ 0.05 (*), p ≤ 0.01 (**), p ≤ 0.0001 (****).

An even clearer picture emerged from the released cytokines in the culture supernatants of this restimulation assay. Splenocytes from infected mice released large amounts of IL-17A (mean 18,345 pg/mL) and IFN-γ (mean 13,234 pg/mL), suggesting re-activation of *S. aureus*-specific Th17 and Th1 cells, respectively (Fig 8, S4 Fig). Moreover, moderate levels of type 2 (IL-13, mean 616.3 pg/mL) and type 4 (IL-10, mean 389.4 pg/mL) cytokines were observed upon re-stimulation. In contrast, splenocytes from PGA-negative control mice released no T cell effector cytokines. A similar, IL-17-dominated pattern was observed in cells from the iliac lymph nodes, which drain the genital area [37]. Mice with PGA also had significantly lower lymphocyte counts than non-infected mice (S2 Table). In summary, these flow cytometry and cytokine data indicate a systemic, Th17-dominated *S. aureus*-specific T cell response in PGA, with less pronounced Th1, Th2, and Treg responses.

**Fig 8.**
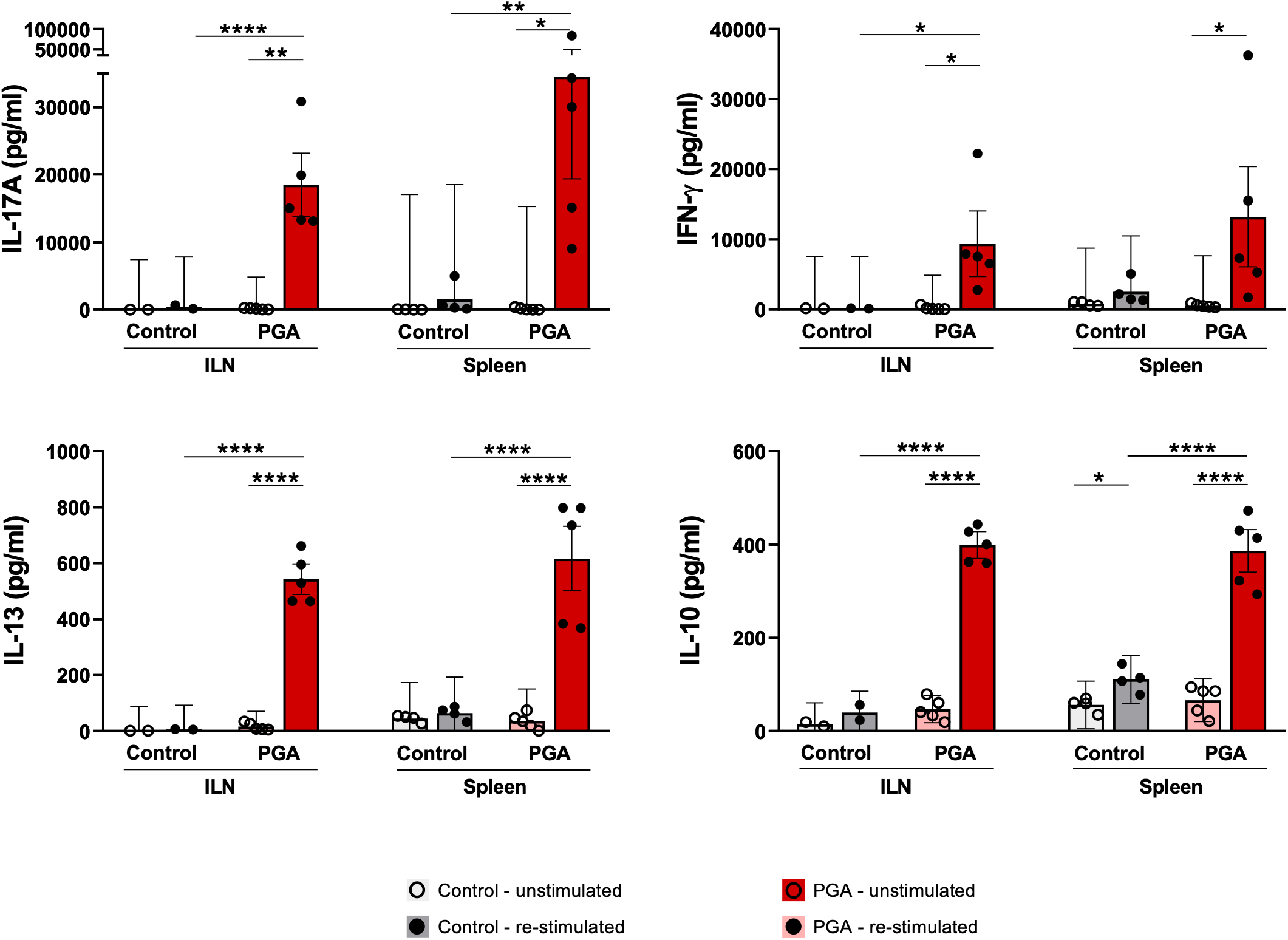
*S. aureus*-stimulated lymph node cells and splenocytes release predominantly Th1- and Th17-related cytokines. Cells from iliac lymph nodes (ILN) or splenocytes from JSNZ-colonized C57BL/6NRj mice with PGA were cultured with or without an *S. aureus* antigen cocktail for 4 days. *S. aureus* CC15 colonized, PGA-negative animals served as control. Cytokines were quantified by bead-based multiplex assay. Each data point represents one mouse (n = 4-5/group), except for ILN data (no PGA group) where one of the two data points presents pooled cells from three mice. Further analyzed cytokines are depicted in S4 Fig. Median ± IQR. Statistics: One-way ANOVA with Sidak correction. p ≤ 0.05 (*), p ≤ 0.01 (**), p ≤ 0.001 (***), p ≤ 0.0001 (****).

### PGA elicits a robust anti-*S. aureus* antibody response

To evaluate the humoral immune response against *S. aureus* JSNZ, we measured serum IgG against UV-killed JSNZΔ*spa* cells by ELISA. Male mice with PGA mounted a strong systemic IgG response (median OD 0.31), whereas colonized mice without PGA had more variable and overall lower IgG levels (median OD 0.09). Naïve *S. aureus*-free mice lacked *S. aureus*-specific antibodies (median OD 0.01), consistent with no prior exposure (Fig 9).

**Fig 9.**
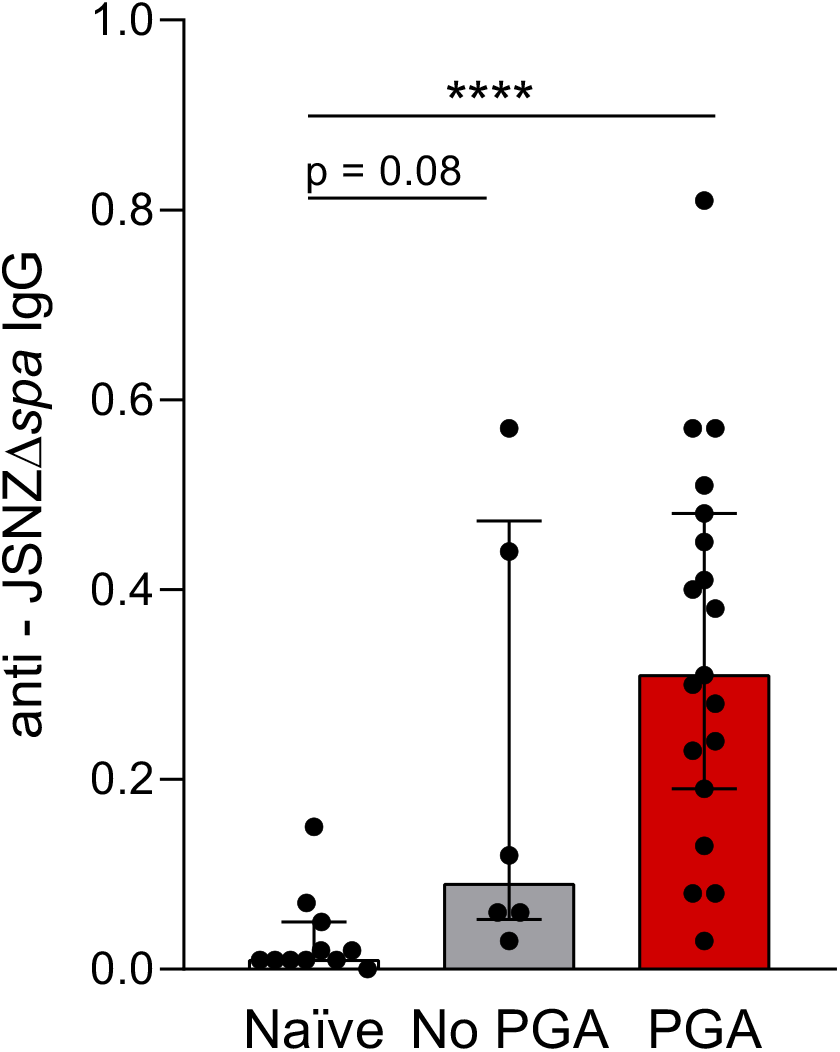
PGA induces a strong systemic anti-JSNZ IgG response. Anti-*S. aureus* serum IgG (1:1,000 dilution) was quantified in naïve (n = 11), No PGA (n = 6), and PGA (n = 19) animals by ELISA using UV-killed JSNZΔ*spa* cells (5 × 10^7^ CFU/mL) as antigen. Each data point represents one mouse (mean of technical duplicates). Median ± interquartile range. Statistics: Kruskal–Wallis test (*p* ≤ 0.0001), with Dunn’s post hoc test. p ≤ 0.0001 (****); ns, not significant.

## Discussion

*S. aureus* infections typically originate from asymptomatic colonization of mucosal surfaces, an aspect underrepresented in existing animal models. Here, we report that persistently colonized male C57BL/6J mice developed spontaneous infections of their preputial glands caused by the colonizing *S. aureus* strain. The resulting abscesses triggered a local inflammation, strong pus formation, and gland destruction. Immune profiling revealed locally elevated cytokines and chemokines, together with a robust systemic antibody and a Th17-dominated T cell response against the invading strain.

The colonizing *S. aureus* strain JSNZ was the causative agent in all PGA cases in our *S. aureus*-positive C57BL/6J breeding colony. This mouse-adapted strain was first isolated during a PGA outbreak in a C57BL/6N colony in 2008 [30]. It was not transmitted from animal staff but likely had an endogenous origin, since many mice in the colony carried it without developing disease [30]. Indeed, mouse-adapted CC88 isolates have circulated among different laboratory mouse breeders for many decades [38,39]. Phage typing of *S. aureus* isolates from a 1967 outbreak of bulbourethral gland infections in a mouse breeding facility likewise indicated an endogenous origin [40]. Notably, these spontaneous infections recapitulate the human situation: Nasal carriage is a well-known risk factor for *S. aureus* infection in both the hospital and community settings, with individuals often infected by their own strain [10,41]. Our colonization-driven infection model now enables the study of specific triggers that disrupt the *S. aureus*–host balance and promote infection [17].

The reasons for the preferred invasion of the preputial glands as compared to the skin or other skin appendages are still unclear [35,42]. Preputial glands are sebaceous glands, containing sebum, pheromonal lipids, phospholipids, wax esters, and triglycerides [43]. *In vitro*, both *S. aureus* and *S. epidermidis* grow robustly at high concentrations of artificial sebum in nutrient rich medium [44]. Moreover, the keratinized, lipid-rich excretory duct supports colonization by certain microbes, including staphylococci [45,46]. This exposed, stratified squamous epithelium could provide a niche for *S. aureus* adherence and persistence, with PGA likely arising from subsequent ascending invasion from the excretory duct. Accordingly, PGA can be experimentally induced by direct inoculation of *S. aureus* into the penile urethra [47], which lies close to where the preputial glands open onto the skin at the prepuce–penis junction [34]. Similar ascending mechanisms occur in otitis externa [48] and mammary duct infections [49], where superficial skin colonization precedes ductal invasion.

Furthermore, the fact that PGA only occurred in sexually mature males suggests a role for male sex hormones. Male mice display elevated androgen production not only in sex organs but also in the skin, modulating tissue-specific gene expression, immune responses and microbial colonization [50– 52]. This aligns with epidemiological data showing higher rates of *S. aureus* SSTIs and nasal carriage in men than women [25,53,54]. Interestingly, testosterone directly activates the Agr quorum-sensing system, a major virulence regulator in *S. aureus* SSTI [55–57]. Thus, high testosterone levels in the preputial gland might switch invading *S. aureus* bacteria to a hypervirulent phenotype, promoting cytotoxicity and abscess formation [56].

In our colony, PGA was restricted to adult males, with no clinical signs in females. Consistently, related purulent infections in females, such as clitoral gland abscesses [58] or periurethral swelling [59], are only anecdotally reported, often with *S. aureus* isolated. Anatomically, clitoral glands are much smaller, with male preputial glands weighing nearly ten times more on average [60,61], so female infections may go unnoticed during routine assessments. More systematic surveillance is therefore warranted to clarify the incidence of endogenous SSTIs in both sexes.

Notably, 67% of colonized breeding males and 68% of the male offspring developed PGA, far exceeding prior reports. For comparison, a large survey of 17,722 mice of undefined *S. aureus* carrier status reported an incidence of just 1.8% [47], and other studies reported 0.4% [62] to 3% [63,64]. Several factors likely explain this disparity. Most importantly, all our animals were *S. aureus*-positive, whereas colonization in commercial SPF facilities ranges from 0% to 20.9% (mean: 6.0%)[31]. As *S. aureus* seems to be the main causative agent, the likelihood for developing PGA is about 17 times higher in our cohort. Secondly, we report prevalence for males only, whereas the comparator studies included both sexes [62–64]. Thirdly, PGA may be overlooked during routine assessments—we observed several palpable but not yet visible cases. Finally, incidence may depend on the mouse strain [65], microbiome, or environmental stressors. The high prevalence of PGA in our *S. aureus*-positive colony thus highlights the real burden of *S. aureus*-associated SSTIs in colonized rodent colonies.

*S. aureus* is the main causative agent of PGA. In the aforementioned cohort of over 17,700 mice, it was isolated from 84% of PGA samples [47]. However, whether all murine *S. aureus* isolates have the capacity to induce PGA remains unclear. We colonized our C57BL/6N animals with JSNZ (spa type t729, CC88), because this is the dominant *S. aureus* lineage in laboratory mice and caused most reported murine PGA cases [30,31]. During the PGA outbreak in New Zealand, which led to the discovery of JSNZ, 8 out of 79 PGA samples were spa typed; all belonged to spa type t729 (CC88) [30]. In another study, 10/13 PGA isolates were CC88 and the remainder CC188 [31]. Similarly, a CC88 isolate (spa type t186) was obtained during a PGA outbreak at Washington University [66]. Broader surveillance and phylogenetic analyses are needed to establish whether additional mouse-adapted *S. aureus* lineages possess similar characteristics that facilitate colonization-driven infection. Besides *S. aureus*, few pathogens have been implicated in natural preputial gland lesions, including *Enterobacter cloacae, Pasteurella pneumotropica, Klebsiella oxytoca, Escherichia coli*, and *Corynebacterium* species [45,47,65,67,68].

PGA presented with a robust but localized inflammatory response. Cytokine profiling of infected preputial gland homogenates revealed marked upregulation of TNF-α, IL-6, IL-17A, IL-23, IL-1α, IL-1β, and the neutrophil-attracting chemokines KC (CXCL1) and MIP-1α. This profile mirrors the Th17-driven type 3 immune response characteristic of *S. aureus* SSTIs [26,69,70]. IL-1α, IL-1β, IL-6, and IL-23 promote IL-17A production by γδ T cells and Th17 cells in skin [70,71], while IL-17A and IL-17F induce chemokines like KC and CXCL2 that drive neutrophil recruitment and activation [71,72,70]. The neutrophilic infiltration observed by immunohistochemistry further confirmed type 3 inflammation. Importantly, IL-17A and IL-22 are essential for bacterial clearance but can delay cutaneous wound healing, indicating a dual role in bacterial clearance and organ pathology [73,74].

A hallmark of PGA was pus accumulation in the preputial gland lumen. Pus consists mainly of dead neutrophils and proteinaceous debris, including neutrophil extracellular traps (NETs)[36]. The colocalization of *S. aureus* clusters with neutrophils in our samples suggests NET formation in the lumen [75]. Although NETs contribute to bacterial containment, their persistence can exacerbate tissue damage and inflammation [75,76]. Overall, the cytokine and chemokine landscape and IHC data reflect a localized, *S. aureus*-specific type 3 immune response aimed at neutrophil mobilization and bacterial containment. Notably, this inflammatory response remained highly localized: elevated cytokine and chemokine levels were detected only in the infected gland, but not in the neighboring healthy gland.

Spontaneous PGA was associated with a robust, antigen-specific Th17-dominated response both in the draining iliac lymph nodes and the spleen. This was evidenced by the proliferation of effector memory CD4^+^RORγt^+^ T cells and marked secretion of IL-17A, IL-17F, and IL-22 after *in vitro* restimulation of splenocytes with *S. aureus* antigens. This systemic Th17-driven response was evident in JSNZ-colonized and – infected males (PGA group), but not in the control mice that were naturally colonized with a CC15 *S. aureus* isolate. In a parallel study, we observed that female mice that were colonized with JSNZ but remained free of symptoms mounted a local and systemic JSNZ-specific Th17-dominated response [33]. This discrepancy may reflect sex differences in antibacterial immunity [25,77], differences in the virulence of JSNZ and the CC15 isolate [38], and low crossreactivity between the JSNZ antigen cocktail and the CC15 isolate.

The Th17 polarization in our mouse abscess model is consistent with prior work implicating Th17 cells and IL-17 in abscess formation and neutrophil recruitment during *S. aureus* infection [69,71,78]. Interestingly, we also observed strong IFN-γ induction and modest increases in IL-13, IL-5, IL-10, and IL-9 in re-stimulated lymph node cells and splenocytes, indicating a mixed cytokine milieu. This is consistent with previous reports of concurrent Th1 and Th2 activation during *S. aureus* infection [20,79]. The anti-inflammatory cytokine IL-10 may counterbalance inflammation and limit tissue damage, but can also facilitate bacterial persistence in localized *S. aureus* infections [21,79,80].

JSNZ colonization induced a low-titer *S. aureus*-specific antibody response, which was strongly increased in animals with PGA. As short-term experimental nasal colonization of human volunteers did not strongly boost anti-staphylococcal antibodies [81], we propose that minor, unnoticed invasive events in colonized mice triggered the low-titer anti-staphylococcal IgG [3]. In contrast, the profound and chronic infection of the preputial glands drove a robust serum antibody response. A similar pattern, higher antibody levels in *S. aureus* infection vs. healthy controls, has been observed in several other studies [82,83]. The JSNZ-specific antibody and Th17 responses, together with neutrophil recruitment, contained but did not clear the infection. This recapitulates the clinical situation, in which SSTIs often last for weeks and recur after antibiotic treatment [5,84]. This scenario is a hallmark of *S. aureus* infections, which do not lead to lasting immune protection.

A limitation of our study is the reliance on visible pathology to estimate prevalence; clinical observation alone likely underestimates the true burden, as subclinical or subtle disease may go unnoticed. While humans lack preputial glands [34], PGA shares many features with *S. aureus* skin- and soft tissue abscesses, the most common manifestation of *S. aureus* infection. Like these abscesses, PGA involves ascending infection of a skin gland through its duct (e.g., sebaceous glands in hair follicles), massive bacterial proliferation, neutrophil infiltration, NET formation, and pus. Moreover, in both settings, the immune response fails to eliminate bacteria or heal the lesion, so surgical intervention or abscess rupture is required.

Overall, PGA is a frequent complication of *S. aureus* colonization in male mice, inducing local inflammation, a strong local and systemic *S. aureus*-specific Th17 response, pronounced neutrophil recruitment, and pus formation. Nevertheless, the animals could not clear the infection. This mirrors the human situation, where *S. aureus* infections often become chronic. Future studies could employ this new model to identify correlates of protection and test interventions for staphylococcal SSTI. The work also has implications for animal welfare and experimentation: *S. aureus* colonization and PGA should be closely monitored in laboratory mouse colonies. Colonized but especially PGA-affected animals mount a strong *S. aureus*-specific adaptive immune response. Thus, unnoticed exposure will affect the outcome of *S. aureus* infection and vaccination studies [85–88]. Moreover, PGA may impair male breeding performance [30]. These findings underscore the need for active monitoring of *S. aureus* colonization and pathology in mouse colonies, especially when immune or reproductive phenotypes are endpoints.

## Methods

### *S.aureus* strain and cultivation

Murine colonization experiments employed the mouse-adapted *S. aureus* strain JSNZ (ST88/CC88-MSSA). JSNZ was initially isolated during an outbreak of PGA among C57BL/6J male mice at the University of Auckland animal facility [30]. A protein A-deficient mutant (JSNZΔ*spa*) was used for *in vitro* re-stimulation assays and antibody profiling [31].

For intranasal inoculation, JSNZ cultures were grown in Brain Heart Infusion (BHI) medium (SigmaAldrich Chemie GmbH, Steinheim, Germany) at 37 °C with shaking to mid-logarithmic growth phase as previously described [32]. For UV-killed *S. aureus* cells, JSNZΔ*spa* was grown in Tryptic soy broth (TSB) (Oxoid Ltd, Hampshire, United Kingdom) from an overnight culture to mid-logarithmic growth phase. Cells were harvested by centrifugation (10,000 *×* g, 10 min, 4°C), washed once with phosphate-buffered saline (PBS, Pan Biotech, Aidenbach, Germany), adjusted to OD (600 nm) = 0.2 with PBS, and exposed to UV light in open petri dishes for 1 hour. After another washing step, cells were adjusted to OD (600 nm) = 0.21 (corresponding to 1×10^8^ CFU/mL) and stored at -80 °C.

### Animal procurement and husbandry

#### *S. aureus*-free mice

Male C57BL/6NRj mice (10 weeks old) with Specific and Opportunistic Pathogen-Free (SOPF) status (i.e., free of *S. aureus*), were obtained from Janvier Labs (Saint-Berthevin, France) and housed at the Central Service and Research Facility for Laboratory Animals at the University Medicine Greifswald, Germany (ZSFV). Animals underwent a seven-day acclimatization period upon arrival. Male control mice (C57BL/6N) obtained for T cell response analysis were obtained from a separate barrier within the ZSFV. These mice were retrospectively found to harbor a CC15 MSSA isolate (spa type t084) in their nasal cavity, a lineage previously detected in our facility [39].

#### S. aureus JSNZ-colonized mice

A *S. aureus*-colonized C57BL/6NRj mouse colony was established at the University Medicine Greifswald under biosafety level 2 (S2) conditions, as previously described [33]. Briefly, male and female mice (see above) were intranasally inoculated with 10^8^ CFU JSNZ and mated seven days later. The offspring were weaned 21 days after birth, and the persistence of *S. aureus* colonization was monitored by quantifying bacterial loads in fecal samples.

#### Housing and Care

Both *S. aureus*-free mice (ZSFV) and *S. aureus*-colonized mice (S2 facility) were housed in individually ventilated cages (IVC, maximum four animals per cage) under SOPF conditions, including autoclaved bedding, enrichment and nesting materials, and ad libitum access to sterilized food and water.

### Determination of the bacterial load

Mouse organs (nose, cecum, lung, heart, spleen, liver, kidneys, and preputial glands) were excised, weighed, and transferred to sterile homogenizer tubes containing zirconium oxide beads (1.4/2.8 mm, Bertin technologies, Precellys, Paris, France). The nose and preputial glands were homogenized in 1 mL of sterile, cold phosphate-buffered saline (PBS) supplemented with a cOmplete™Mini, EDTA-free Protease inhibitor cocktail (Roche, Basel, Switzerland) at 6,500 rpm for 2 × 30 s (5 min interval) using the Precellys 24 lysis and homogenization system (Bertin Technologies). All other organs were homogenized in 1 mL of sterile, cold PBS at 6,000 rpm for 2 × 20 s (15 s interval).

Fecal specimens were suspended in sterile cold PBS at 0.2 g/mL and homogenized at 1,400 rpm for 20 min at 4°C using a Thermo Mixer C shaker (Eppendorf, Hamburg, Germany). Serial dilutions of all homogenates were plated in triplicate (10 µL) on *S. aureus* CHROMagar (CHROMagar, Paris, France). and incubated at 37°C for 16 h. Colonies were enumerated by standard plate counting and expressed as colony-forming units (CFU) per organ or CFU/mL for fecal samples.

### Microbiological identification of *S. aureus* and other species

*S. aureus* was identified in preputial gland homogenates by colony morphology on mannitol salt agar (MSA) plates and confirmed by a latex agglutination test (Pastorex Staph Plus, Bio-Rad Laboratories GmbH, Feldkirchen, Germany. Non-*S. aureus* species were identified by 16S ribosomal RNA (rRNA) gene sequencing using primers 1_27F (5⍰-aga gtt tga tcm tgg ctc ag-3⍰) and 16S_2 (5⍰-ccg tca att cmt ttg agt tt-3⍰). PCR products were purified (QIAquick, QIAGEN GmbH, Hilden, Germany) and Sanger sequenced (Eurofins MWG, Ebersberg, Germany). Reads were trimmed and aligned using MUSCLE. Species identification was performed via BLASTn against NCBI reference databases.

### Spa genotyping

Spa genotyping of *S. aureus* strains was performed according to published protocols using the primers spa-1113f and spa-1514r [89]. *Spa* types were assigned to clonal complexes using Ridom SpaServer (http://www.spaserver.ridom.de).

### Paraffin embedding and Hematoxylin and Eosin (H&E) staining

Preputial glands were fixed overnight in 37% formaldehyde (Carl Roth, Karlsruhe, Germany), dehydrated through a graded ethanol series (70%, 80%, 96%, 100% x2; 1h each), cleared in 100% xylene (J.T. Baker, Avantor, Deventer, Netherlands), infiltrated with molten paraffin, and embedded in paraffin wax (Süsse Labortechnik GmbH). Sections of 3–5 μm were cut using a rotary microtome (Leica RM2255, Leica Biosystems, Wetzlar, Germany).

For H&E staining, sections were deparaffinized in 100% xylene (J.T. Baker), rehydrated through a descending ethanol series (100%, 96%, 80%, and 70%, 5 min each step), and rinsed with distilled water. Sections were stained with alum haematoxylin (Sigma Aldrich Chemie GmbH) for 3 min 30 s at RT, rinsed in distilled water, and blued in tap water. Counterstaining was performed with Eosin Y (Epredia, Microm International GmbH, Dreieich, Germany) for 20 s. Sections were dehydrated through an ascending ethanol series, cleared in xylene, and mounted with VitroClud (R. Langenbrinck GmbH, Emmendingen, Germany). Bright-field imaging was performed with a hybrid microscope (REB-01-D, ECHO, California, United States) equipped with a 12-megapixel CMOS color camera, using ECHO Pro software (v5.3.1) with calibrated scale bars.

### Immunohistochemistry (IHC)

IHC was performed as previously described [90]. Formalin-fixed, paraffin-embedded (FFPE) preputial gland sections (4 μm) were deparaffinized in xylene and rehydrated through a graded ethanol series. Antigen retrieval was carried out using heat-induced epitope retrieval (HIER) (ScyTek Laboratories, Utah, USA) in 10% glycerol buffer (Carl Roth, Karlsruhe, Germany) at 70°C for 120 min. After rinsing with deionized water and Tris-buffered saline (TBS; pH 7.4), sections were blocked for 30 min at RT in a blocking solution containing 1% bovine serum albumin (BSA), 2% normal donkey serum (Millipore, Merck KGaA, Darmstadt, Germany), 5% cold water fish gelatin, 0.05% Tween-20, and 0.05% Triton X-100 (all Sigma Aldrich Chemie GmbH).

Sections were incubated with primary antibodies against murine neutrophils (monoclonal rat anti-GR1, RB6-8C5, 1:50; eBioscience GmbH, Frankfurt a. Main, Germany) and *S. aureus* (polyclonal rabbit anti-*S. aureus*-FITC, 1:30; Invitrogen GmbH, Darmstadt, Germany) for 90 min at RT. This was followed by incubation with species-specific secondary antibodies (polyclonal rabbit anti-rat, Alexa 647, 1:200; polyclonal rabbit anti-fluorescein, Alexa 488; Invitrogen GmbH, Darmstadt, Germany). Nuclei were counterstained with DAPI (1:100 in PBS; Molecular Probes, Thermo Fisher Scientific, Dreieich, Germany) for 1 min, and sections were mounted using Dako Fluorescent Mounting Medium (Agilent Technologies, Waldbronn, Germany). Imaging was performed using a Keyence BZ-9000 fluorescence microscope (Keyance, Neu-Isenburg, Germany).

### Peroxidase activity assay

50 µL of homogenized preputial glands were diluted 1:10 or 1:100 in PBS and incubated with 150 µl of a peroxidase chromogenic substrate (BD OptEIA™ TMB substrate, BD Biosciences) for 15 min. The reaction was stopped with 50 µl of 2N H_2_SO_4_ and absorbance was measured at 450 nm (reference wavelength: 500 nm) using a microplate reader (TECAN Infinite M200 Pro, Tecan, Männedorf, Switzerland).

### Detection of cytokines and chemokines

Cytokine and chemokine levels in murine serum, preputial gland homogenates and cell-free culture supernatants (all at 1:2 dilution) were quantified using three bead-based multiplex immunoassays (all BioLegend GmbH, Koblenz, Germany): The LEGENDplex™ Mouse Inflammation Panel (13-plex; IL-1α, IL-1β, IL-6, IL-10, IL-12p70, IL-17A, IL-23, IL-27, CCL2 (MCP-1), IFN-β, IFN-γ, TNF-α, and GM-CSF), the Th Cytokine Panel (12-plex; IL-2, IL-4, IL-5, IL-6, IL-9, IL-10, IL-13, IL-17A, IL-17F, IL-22, IFN-γ, and TNF-α), and the Mouse Proinflammatory Chemokine Panel (13-plex, MCP-1 (CCL2), RANTES (CCL5), IP-10 (CXCL10), Eotaxin (CCL11), TARC (CCL17), MIP-1α (CCL3), MIP-1β (CCL4), MIG (CXCL9), MIP-3^α^ (CCL20), LIX (CXCL5), KC (CXCL1), BLC (CXCL13), and MDC (CCL22)). Samples were processed per manufacturer’s instructions, and measured on an LSR II flow cytometer (BD Biosciences, Heidelberg, Germany). Data were analyzed using the LEGENDplex™ software Qognit. Analytes below the limit of detection (LOD) were set to LOD/2.

### Quantification of anti-*S. aureus* JSNZ IgG

Serum IgG against *S. aureus* were measured by ELISA using UV-killed *S. aureus* JSNZΔspa cells as coating antigen. Plates were coated with 5 × 10^7^ CFU/mL in 1× Candor buffer (pH 9.6; Candor Bioscience, Bayern, Germany) overnight at 4°C, washed three times with PBS + 0.05% Tween-20, and blocked with 100 μL of 10% fetal bovine serum in PBS for 1 h at RT (100 rpm). Serum samples diluted 1:1,000 in blocking buffer were added (50 μL/well) and incubated for 1 h at RT (100 rpm). Plates were washed thrice and incubated with a polyclonal goat anti-mouse IgG, POD-labelled, secondary antibody (Jackson ImmunoResearch Europe Ltd, Cambridgeshire, United Kingdom, 1:20,000) for 1 h at RT with shaking. Plates were washed and developed with BD OptEIA™ TMB substrate (BD Biosciences) for 15 min in the dark at RT. The reaction was stopped with 20 μL of 2N H_2_SO_4_, and absorbance was measured at 450 nm using a microplate reader (TECAN Infinite M200 Pro, Tecan, Männedorf, Switzerland).

### Haematological Analysis

Whole blood collected in EDTA-coated tubes was stored on ice and analyzed within 3 h on a VetScan HM5™ automated haematology analyzer (ABAXIS Europe GmbH, Griesheim, Germany), performing a three-part differential count. The parameters measured included total WBC, lymphocytes, monocytes, neutrophils, RBC, hemoglobin, hematocrit, MCV, platelets, MPV, MCH, and MCHC.

### Preparation of single cell suspensions from tissues

Iliac lymph nodes (ILN) were meshed through a 70 µm cell strainer, washed with 25 mL 5% FBS/PBS, centrifuged (300 × g, 8 min, 4°C), and resuspended in 400 µl of 5% FBS/PBS. Splenocytes were isolated as previously reported [33]. Cell concentrations were determined on an LSR II using Trucount™ beads (BD Biosciences).

### CFSE labelling

Splenocytes and ILN cells were labeled with Carboxyfluorescein diacetate succinimidyl ester (CFDASE) (BioLegend GmbH, Koblenz, Germany) as previously described [33]. Briefly, 4 × 10^6^ cells/mL in PBS were incubated with 4 µM CFDA-SE for 5 min at 37°C in the dark, then washed with 40 mL of 10% FBS/PBS (300 × g, 8 min, RT), and once with T10F medium (TexMACS (Miltenyi Biotec, Bergisch Gladbach, Germany) containing 10% FBS, 1× Penicillin/Streptomycin/Gentamycin (Gibco Invitrogen, Carlsbad, USA), and 50 µM β-mercaptoethanol (Merck, Darmstadt, Germany)). Labelled cells were resuspended in T10F at 1 × 10^7^ cells/mL.

### In vitro re-stimulation with S. aureus antigens

Cells were left unstimulated or stimulated with an *S. aureus* antigen cocktail consisting of heatinactivated extracellular proteins (5 µg/ml) combined with UV-inactivated cells of JSNZΔ*spa* (2.5 × 10^5^ CFU/ml) for 4 days at 37°C and 5% CO_2_. Cell-free culture supernatants were stored at -80°C for cytokine analysis.

### Flow cytometry

CFSE-labeled cells were washed twice with PBS (300 × g, 8 min, 4°C) and stained with Zombie NIR™ (BioLegend GmbH) for 30 min at RT in the dark. After washing with 2 ml FACS buffer (2% FBS, 0.02% Sodium Azide (NaN_3_), 2 mM EDTA in PBS), cells were incubated with FcR block solution (Miltenyi Biotec) for 5 min at 4 °C. Next, the cells were surface-stained for 20 min at 4°C with antibodies specific to CD3 (REA641, PerCP-Vio 700), CD4 (REA604, Vio R720), CD8 (REA601, VioGreen), CD62L (MEL-14, BV605, Biolegend) and CD44 (REA664, Pe-Vio770) (all Miltenyi Biotec, except CD62: BioLegend) (S1 Table). For intranuclear transcription factor staining, cells were fixed and permeabilized using the MACS Transcription Factor Buffer Set (Miltenyi Biotec) per manufacturer’s protocol, and then stained intracellularly for 30 min at 4°C with antibodies specific to RORγt (REA278, PE), FOXP3 (REA788, VioR667), GATA3 (REA174, PE-Vio615; all Miltenyi Biotec), and T-bet (04-46, BV650, BD Biosciences). Fluorescence Minus One (FMO) controls were used for transcription factor gating. Cells were resuspended in 200 µl FACS buffer containing BD Trucount™ beads. Flow cytometric data were acquired on an LSR II and were analyzed using the software FlowJo™ V10 (BD BioSciences). Gating strategy is displayed in S5 Fig.

### Ethics statement

All animal experiments were approved by the local State authorities (Landesamt für Landwirtschaft, Lebensmittelsicherheit und Fischerei Mecklenburg-Vorpommern, approval no, 7221.3-1-058/20) and conducted in accordance with the German Animal Welfare Act (Deutsches Tierschutzgesetz) and the EU Directive 2010/63/EU.

### Statistical Analysis and data visualization

Statistical analyses and data visualization were performed in GraphPad Prism 8 package (GraphPad Software, Inc., La Jolla, United States) and R (core v.4.5.0) [91] for statistical computing (tidyverse package v.2.0.0 [92]; rstatix package v.0.7.2 [93]) and data visualization (ggplot2 package v3.5.2.) [94]. To compare two datasets, we employed a Mann-Whitney test. To compare multiple groups, we employed a Kruskal-Wallis test with Dunn’s multiple comparison test. Data from re-stimulation assays present a combination of paired and unpaired samples and were analysed by a linear mixed model in two-way ANOVA design. Details of statistical analyses, including statistical tests and significance criteria, are provided in the figure legends.

## Supplementary Material

**S1 Fig. *S. aureus* does not disseminate systemically in mice with PGA**. *S. aureus* CFUs were quantified in the lungs, liver, kidneys, heart, spleen, and urine using CHROMagar selective plates. Groups include: naïve (*S. aureus*-free mice), No PGA (JSNZ-colonized mice without preputial gland infection), and PGA (JSNZ-colonized mice with preputial gland infection). Each data point represents one animal (CFU value is the median of three technical replicates). Median ± IQR. ND, not determined.

**S2 Fig. Serum cytokines are not affected by PGA**. Heat map illustrating log_10_-transformed and hierarchically clustered serum cytokine and chemokine concentrations. Groups: naïve (*S. aureus*-free); no PGA (JSNZ-colonized, no PGA); PGA (*S. aureus* JSNZ-colonized, with PGA). For IL-2 and IL-17F, all samples below the detection limit were set at LOD/2.

**S3 Fig. Absolute cell counts for re-stimulated splenocytes from JSNZ-colonized PGA-positive mice and PGA-free controls**. Splenocytes from JSNZ-colonized mice with PGA were CFSE-labelled and cultured with or without an *S. aureus* antigen cocktail for 4 days. Mice initially considered *S. aureus*-naïve but retrospectively found to carry a CC15 isolate served as the colonized PGA-negative control. CD4^+^ and CD8^+^ T cell proliferation and the frequency of lineage-specific transcription factors RORγt, T-bet, GATA3 and FOXP3 within Th cell populations were assessed by flow cytometry. Within CD4^+^ T cells, effector/effector memory T cells (Teff/Tem, CD44^+^CD62L^-^) and central memory T cells (Tcm, CD44^+^CD62L^+^) subsets were identified. Data are displayed as absolute cell numbers (n = 4-5/group). Mean ± CI. Statistics: linear mixed model (including animal-ID as random factor) in two-way ANOVA design. p ≤ 0.05 (*), p ≤ 0.01 (**), p ≤ 0.0001 (****).

**S4 Fig. *S. aureus*-specific T cells release Th17-, Th1-, Th2-, Treg-, and Th9-related cytokines**. Cells from iliac lymph nodes (ILN) or splenocytes from JSNZ-colonized mice with PGA were cultured with or without an *S. aureus* antigen cocktail for 4 days. *S. aureus* CC15-colonized, PGA-negative animals served as control. Cytokines were quantified by bead-based multiplex assay. Each data point represents one mouse (n = 4-5/group), except for ILN data (no PGA group) where one of the two data points presents pooled cells from three mice. Median ± CI. Statistics: linear mixed model (including animal-ID as random factor) in two-way ANOVA design. p ≤ 0.05 (*), p ≤ 0.01 (**), p ≤ 0.001 (***), p ≤ 0.0001 (****).

**S5 Fig. Flow cytometry gating strategy for identifying murine CD4**^**+**^ **and CD8**^**+**^ **T cell subsets in the spleen**. CFSE-labelled spleen single-cell suspensions were cultured for 4 days in the with or without an *S. aureus* antigen cocktail, then stained for surface and intracellular markers. Singlets were gated using FSC/SSC, live cells were identified by excluding Zombie NIR™ positive cells. Live single cells were gated for leukocytes (SSC-A/FSC-A), followed by selection of T cell population (CD3^+^). CD4^+^, and CD8^+^ T cell subsets were gated within total CD3^+^, and proliferating (CFSE^low^) CD3^+^. Memory subpopulations were defined using CD44 and CD62L: central memory (Tcm, CD44^+^CD62L^+^) and effector memory (Tem, CD44^+^CD62L^-^). Expression of T-bet, RORγt, and FOXP3 was assessed within each T cell subset, proliferating T cells, and proliferating memory T cells. Presented data are from re-stimulated splenocytes from a PGA mouse.

## Acknowledgements

We thank Dr. Daniel Mrochen and Susanne Neumeister for their support in flow cytometry analyses. We also thank Fawaz Alsholui and Sabine Prettin for excellent technical support in microbiological analyses and animal handling, respectively. Amelie Müller contributed to cytokine quantifications. Dirk Stobbe and Antje Müller supported immunohistochemistry analyses.

Portions of this manuscript were edited for language, clarity, and consistency using Claude (Anthropic, San Francisco, CA, USA). All AI-generated suggestions were reviewed and approved by the authors, who take full responsibility for the content.

## Related manuscript

A manuscript describing the persistent, lifelong neonatal colonization induced by JSNZ and the elicited Th17-dominated host immune response is submitted back-to-back to this manuscript to PLoS Pathogens (Fernandes Hartzig et al).

