## Supplemental Figures 1-5 for "Spontaneous preputial gland infection in *Staphylococcus aureus*-colonized male C57BL/6 mice triggers a Th17-driven immune response"

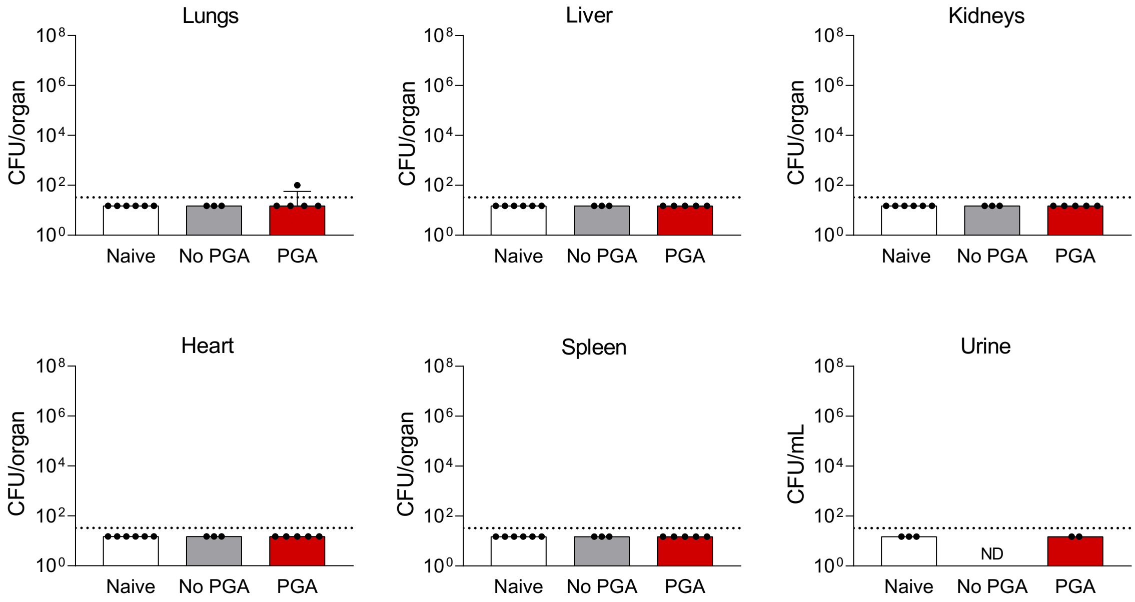


**S1 Figure. *S. aureus* does not disseminate systemically in mice with PGA.** *S. aureus* CFUs were quantified in the lungs, liver, kidneys, heart, spleen, and urine using CHROMagar selective plates. Groups include: naïve (*S. aureus*-free mice), No PGA (JSNZ-colonized mice without preputial gland infection), and PGA (JSNZ-colonized mice with preputial gland infection). Each data point represents one animal (CFU value is the median of three technical replicates). Median ± IQR. ND, not determined.

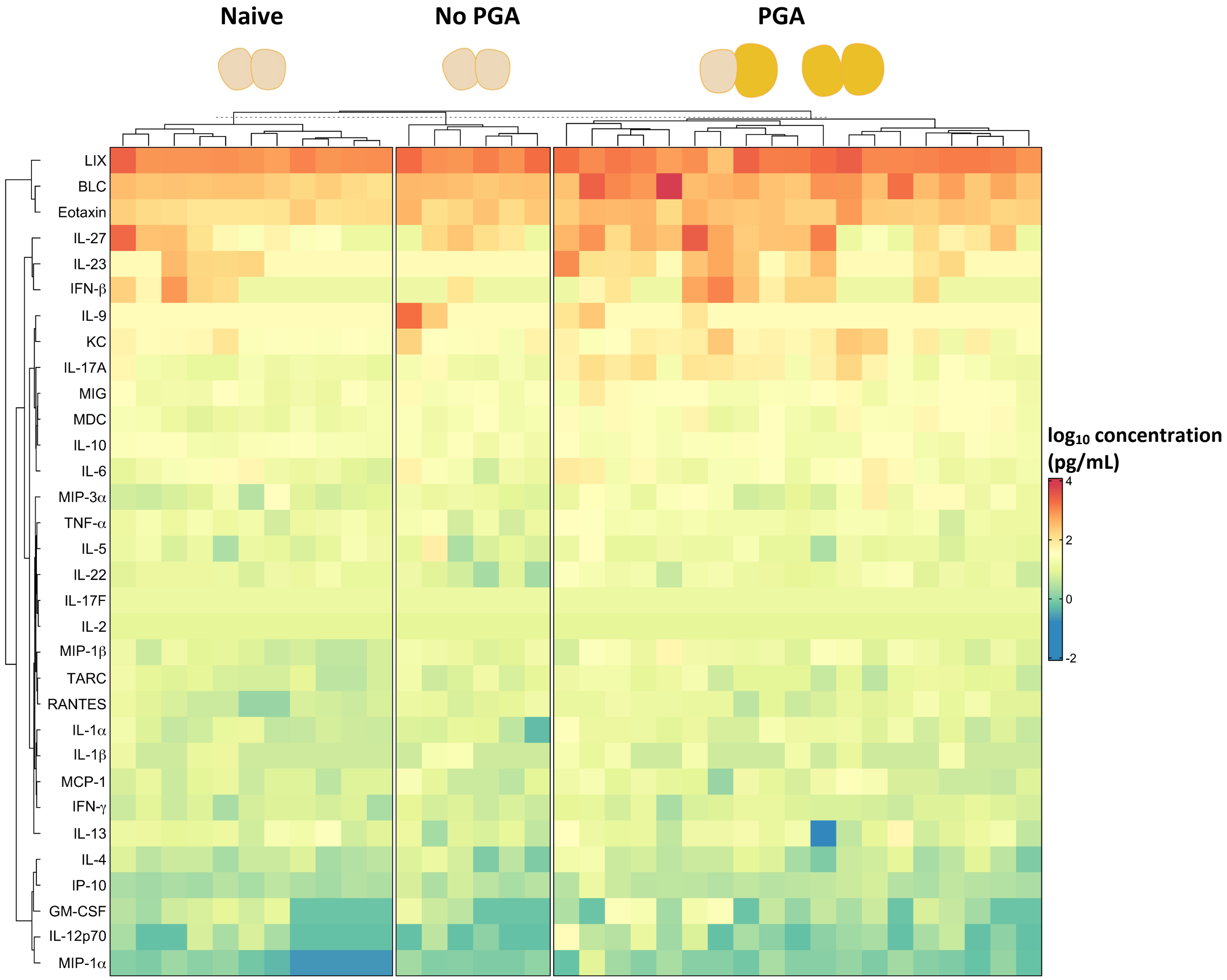


**S2 Figure. Serum cytokines are not affected by PGA.** Heat map illustrating $log_{10}$-transformed and hierarchically clustered serum cytokine and chemokine concentrations. Groups: naïve (*S. aureus*-free); no PGA (JSNZ-colonized, no PGA); PGA (*S. aureus* JSNZ-colonized, with PGA). For IL-2 and IL-17F, all samples below the detection limit were set at LOD/2.

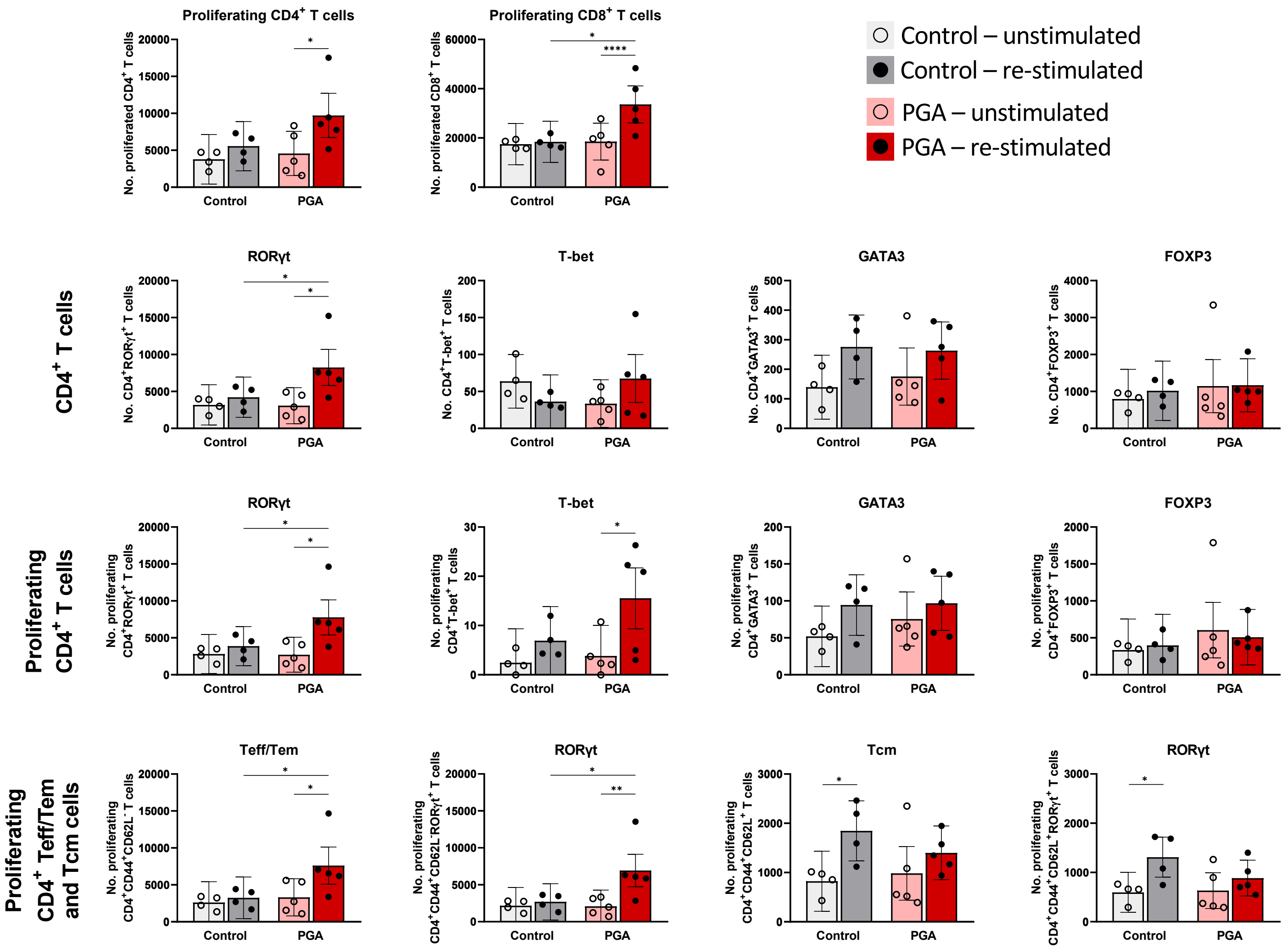


**S3 Figure. Absolute cell counts for re-stimulated splenocytes from JSNZ-colonized PGA-positive mice and PGA-free controls.** Splenocytes from JSNZ-colonized mice with PGA were CFSE-labelled and cultured with or without an *S. aureus* antigen cocktail for 4 days. Mice initially considered *S. aureus*-naïve but retrospectively found to carry a CC15 isolate served as a colonized PGA-negative control. $CD4^+$ and $CD8^+$ T cell proliferation and the frequency of lineage-specific transcription factors RORγt, T-bet, GATA3 and FOXP3 within Th cell populations were assessed by flow cytometry. Within $CD4^+$ T cells, effector/effector memory T cells (Teff/Tem, $CD44^+CD62L^-$) and central memory T cells (Tcm, $CD44^+CD62L^+$) subsets were identified. Data are displayed as absolute cell numbers (n = 4-5/group). Mean ± CI. Statistics: linear mixed model (including animal-ID as random factor) in two-way ANOVA design. p ≤ 0.05 (*), p ≤ 0.01 (**), p ≤ 0.0001 (****).

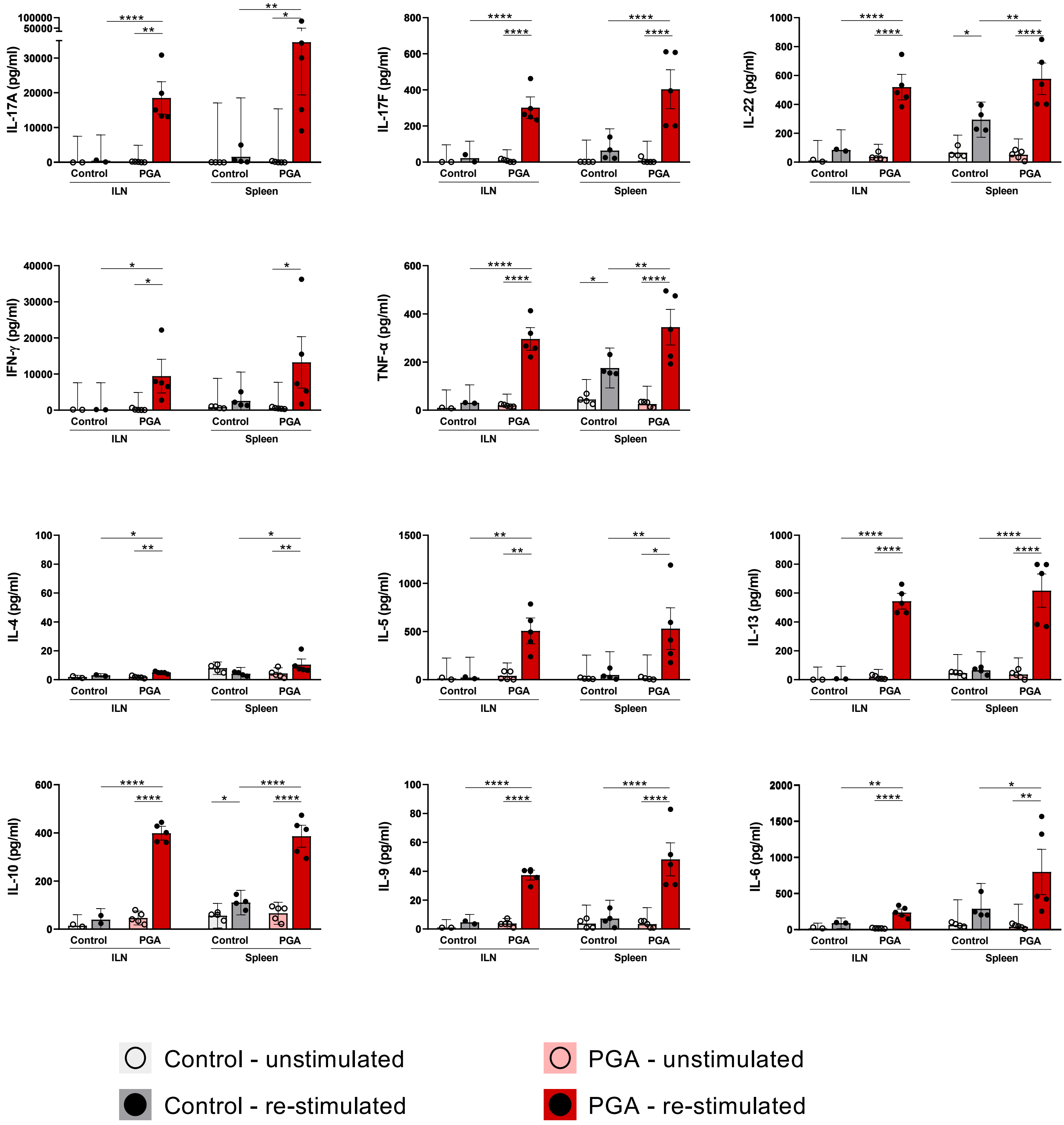


**S4 Figure. *S. aureus*-specific T cells release Th17-, Th1-, Th2-, Treg-, and Th9-related cytokines.** Cells from iliac lymph nodes (ILN) or splenocytes from JSNZ-colonized mice with PGA were cultured with or without an *S. aureus* antigen cocktail for 4 days. *S. aureus* CC15-colonized, PGA-negative animals served as control. Cytokines were quantified by bead-based multiplex assay. Each data point represents one mouse (n = 4 – 5/group), except for ILN data (no PGA group) where one of the two data points presents pooled cells from three mice. Median ± CI. Statistics: linear mixed model (including animal-ID as random factor) in two-way ANOVA design. $p \leq 0.05$ (*), $p \leq 0.01$ (**), $p \leq 0.001$ (***), $p \leq 0.0001$ (****).

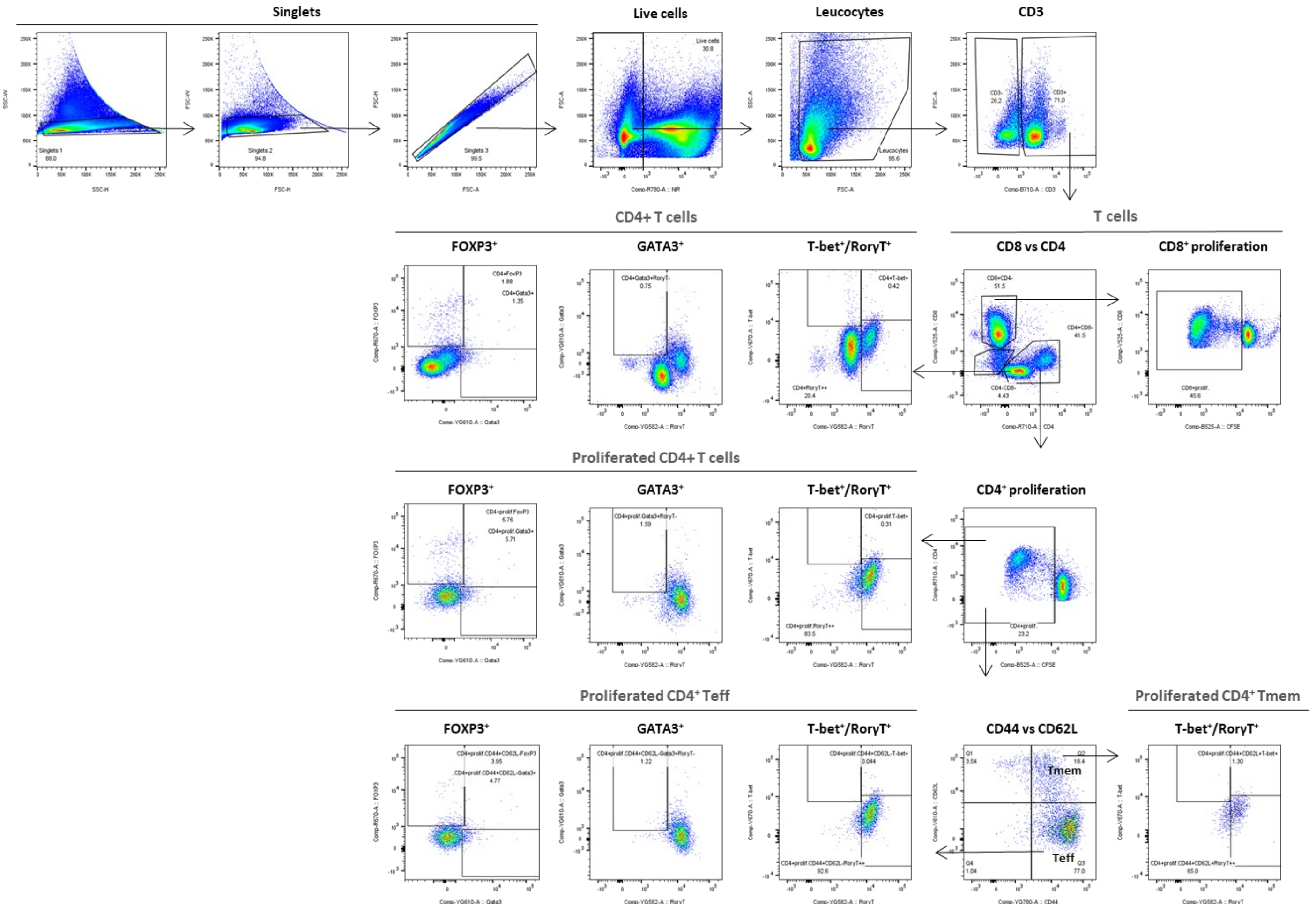


**S5 Figure. Flow cytometry gating strategy for identifying murine CD4$^{+}$ and CD8$^{+}$ T cell subsets in the spleen.** CFSE-labelled spleen single-cell suspensions were cultured for 4 days in the with or without an *S. aureus* antigen cocktail, then stained for surface and intracellular markers. Singlets were gated using FSC/SSC, live cells were identified by excluding Zombie NIR™ positive cells. Live single cells were gated for leukocytes (SSC-A/FSC-A), followed by selection of T cell population (CD3$^{+}$). CD4$^{+}$, and CD8$^{+}$ T cell subsets were gated within total CD3$^{+}$, and proliferating (CFSE$^{low}$) CD3$^{+}$. Memory subpopulations were defined using CD44 and CD62L: central memory (Tcm, CD44$^{+}$CD62L$^{+}$) and effector memory (Tem, CD44$^{+}$CD62L$^{-}$). Expression of T-bet, RORγt, and FOXP3 was assessed within each T cell subset, proliferating T cells, and proliferating memory T cells. Presented data are from re-stimulated splenocytes from a PGA mouse.
