## Supplemental Tables 1-2 for "Spontaneous preputial gland infection in *Staphylococcus aureus*-colonized male C57BL/6 mice triggers a Th17-driven immune response"

**S1 Table. Antibodies used for flow cytometry**

| **Marker** | **Conjugate** | **Channel** | **Clone** | **Isotype** | **Concen-tration**  **(mg/mL)** | **Dilution** | **Company** | **Catalogue No.** |
| --- | --- | --- | --- | --- | --- | --- | --- | --- |
| CD3 | PerCP-Vio700 | B710 | REA641 | REA^1^, human IgG1 | 0.15 | 1:25 | Miltenyi Biotec | 130120826 |
| CD4 | Vio R720 | R710 | REA604 | REA^1^, human IgG1 | 0.15 | 1:50 | Miltenyi Biotec | 130127473 |
| CD8 | VioGreen | V525 | REA601 | REA^1^, human IgG1 | 0.15 | 1:25 | Miltenyi Biotec | 130122017 |
| CD62L | BV605 | V610 | MEL-14 | Rat IgG2a, κ | 0.2 | 1:50 | BioLegend | 104438 |
| CD44 | PE-Vio770 | YG780 | REA664 | REA^1^, human IgG1 | 0.15 | 1:50 | Miltenyi Biotec | 130119127 |
| RORγt | PE | YG582 | REA278 | REA^1^, human IgG1 | 0.15 | 1:25 | Miltenyi Biotec | 130123248 |
| FOXP3 | VioR667 | R670 | REA788 | REA^1^, human IgG1 | 0.15 | 1:50 | Miltenyi Biotec | 130111682 |
| T-bet | BV650 | V670 | 04-46 | Mouse IgG1, κ | 5 µl/test | 1:25 | BD Biosciences | 564142 |
| GATA3 | PE-Vio615 | YG610 | REA174 | REA^1^, human IgG1 | 0.15 | 1:50 | Miltenyi Biotec | 130109161 |

^1^ REAfinity^TM^ monoclonal antibody

**S2 Table. Leukocyte alterations in mice with preputial gland infection**

| **Parameters** | **Units** | **Naïve** | **No PGA** | **PGA** | **C57BL/6J^2^** | **C57BL/6J^2^** | **C57BL/6J^2^** | **p value^1^** |
| --- | --- | --- | --- | --- | --- | --- | --- | --- |
| Age | Weeks | 33.9 ± 3.2 | 22.4 ± 4.0 | 41.1 ± 23.0 | 26* | 52* | 78* |  |
| White Blood Cell count (WBC) | 10^3^ cells/μL | 3.4 ± 0.7 | 3.9 ± 1.1 | 2.5 ± 0.5 | 5.2 ± 1.7 | 5.1 ± 2.4 | 7.2 ± 3.7 | 0.1069 |
| Lymphocytes (LYM) | 10^3^ cells/μL | 2.53 ± 0.61 | 2.59 ± 0.93 | 1.52 ± 0.21 | 4.39 ± 1.47 | 4.15 ± 2.05 | 6.00 ± 2.84 | **0.0412** |
| Monocytes (MON) | 10^3^ cells/μL | 0.24 ± 0.12 | 0.23 ± 0.05 | 0.21 ± 0.07 | 0.25 ± 0.17 | 0.26 ± 0.18 | 0.35± 0.27 | 0.6611 |
| Neutrophils (NEU) | 10^3^ cells/μL | 0.69 ± 0.53 | 1.06 ± 0.42 | 0.82 ± 0.40 | 0.47 ± 0.21 | 0.62 ± 0.31 | 0.81± 0.77 | 0.5123 |
| Red Blood Cell count (RBC) | 10^6^ cells/μL | 9.3 ± 0.3 | 9.6 ± 0.8 | 8.9 ± 1.4 | 8.2 ± 0.6 | 8.1 ± 0.8 | 8.6 ± 1.0 | 0.5785 |
| Haemoglobin (HGB) | g/dL | 13.5 ± 0.3 | 14.1 ± 1.1 | 12.3 ± 1.4 | 11.5 ± 0.8 | 11.2 ± 1.0 | 11.8 ± 1.3 | 0.1655 |
| Haematocrit (HCT) | % | 39.4 ± 1.6 | 40.9 ± 3.9 | 36.5 ± 3.7 | 36.6 ± 2.0 | 34.8 ± 3.5 | 36.9 ± 4.4 | 0.2199 |
| Mean cell volume (MCV) | fL | 42.0 ± 1.0 | 42.6 ± 0.4 | 41.4 ± 3.4 | 44.8 ± 2.4 | 43.1 ± 1.2 | 43.0 ± 1.3 | 0.3952 |
| Platelet count (PLT) | 10^3^ cells/μL | 743 ± 83 | 673 ± 169 | 953 ± 251 | 951 ± 226 | 852 ± 326 | 1168 ±279 | 0.1891 |
| Mean platelet volume (MPV) | fL | 5.8 ± 0.4 | 5.7 ± 0.1 | 5.7 ± 0.5 | 5.5 ± 0.8 | 5.2 ± 0.7 | 5.1 ± 0.3 | 0.9863 |
| Mean cell haemoglobin (MCH) | pg | 14.4 ± 0.4 | 14.7 ± 0.3 | 13.9 ± 0.8 | 14.1 ± 0.5 | 13.9 ± 0.6 | 13.7 ± 0.7 | 0.2332 |
| Mean cell haemoglobin concentration (MCHC) | g/dL | 34.3 ± 1.5 | 34.5 ± 0.7 | 33.6 ± 0.8 | 31.5 ± 1.4 | 32.3 ± 1.4 | 31.9 ± 1.2 | 0.2471 |

^1^ Data are presented as mean ± SD. Statistical comparisons between the No PGA and PGA groups used unpaired two-tailed t-tests.

^2^Reference values from age-matched C57BL/6 male mice (26, 52, and 78 weeks) were procured from the Jackson Laboratory database. https://media.jax.org/m/7f8ff8eb6db658fb/original/aged-b6-physiological-data-summary.pdf
